# A comprehensive, open-source battery of movement imagery ability tests: Development and psychometric properties

**DOI:** 10.1101/2025.10.20.683365

**Authors:** Marcos Moreno-Verdú, Baptiste M Waltzing, Elise E Van Caenegem, Carla Czilczer, Laurine F Boidequin, Charlène Truong, Stephan Frederic Dahm, Robert M Hardwick

**Affiliations:** Brain, Action, and Skill Laboratory (BAS-Lab), Institute of Neuroscience (Cognition and Systems Division), UCLouvain, Belgium; Department of Psychology, Faculty of Psychology and Sports Sciences, University of Innsbruck, Innsbruck, Austria

**Keywords:** movement imagery, motor imagery, psychometrics, validity, reliability, ability

## Abstract

Imagining actions is a covert and multidimensional skill difficult to quantify. Comprehensive assessments rarely combine measures of imagery generation, maintenance, and manipulation. We developed and validated a combination of tests to assess these processes of movement imagery, online. 180 healthy individuals completed the MIQ-RS questionnaire (generation), the Imagined Finger Sequence Task (iFST; maintenance), and the Hand Laterality Judgement Task (HLJT; manipulation). MIQ-RS showed a bifactorial structure (visual and kinesthetic modalities) according to confirmatory factor analysis, and its reliability (internal consistency) was good. In the iFST, internal validity analyses via generalized mixed models showed a clear effect of sequence complexity, stronger for execution than imagery. Reliability, estimated via signal-to-noise ratios (SNRs) using hierarchical Bayesian models, was also adequate (SNR ≥ 1.6). In the HLJT, expected effects of rotation angle, hand view, and their interaction, consistent with biomechanical constraints, were also found. Reliability was also adequate (SNR ≥ 1.75). Criterion validity across tests, assessed using Bayesian Spearman’s correlations, showed that correlations were generally absent (BF_01_ ≥ 3), and when present, of small magnitude (r ≤ 0.27). Test-retest reliability (123 participants reassessed 6-8 days after), computed via Intraclass Correlation Coefficients (ICCs), was generally adequate (ICCs ≥ 0.67). We conclude that the online versions of these tests showed adequate structural/internal validity and (test-retest) reliability. However, weak criterion validity suggests individuals with high ability to generate movement imagery may not necessarily have high ability to maintain and/or manipulate movement imagery, underscoring the need for comprehensive assessment of this capacity.

## INTRODUCTION

Movement imagery can be defined as mentally performing an action without physically executing it ^1^. Repetitive practice with movement imagery has measurable effects on motor performance ^2,3^. Nonetheless, the effects of this practice may depend on the individual’s ability to perform movement imagery ^4^. Existing models agree that movement imagery requires three processes to be completed successfully ^5,6^: 1) to generate the internal representation of the action, 2) to maintain it until the action is completed, and 3) to manipulate it to dynamically update its content as the action evolves.

An unaddressed challenge has been to develop valid and reliable measures of ‘movement imagery ability’ (i.e. the ability to perform movement imagery) ^7^. In fact, a direct, objective and comprehensive measure of imagery ability has not been developed yet. This is partly because of the ‘multidimensional’ nature of movement imagery: different types of measures aim to assess the different cognitive processes that unfold during imagery (generation, maintenance and manipulation). These measures are typically used separately and are very rarely combined in the same individuals. Because these three processes occur simultaneously during movement imagery, assessing all of them is necessary to comprehensively determine the ability to perform it ^8^. While there have been previous attempts to develop ‘indices’ combining behavioural and physiological tools ^6,9^, this practice has not been widely adopted, and did not consider all three mentioned processes. We argue that the combination of multiple tasks to assess the different processes of movement imagery is valuable to provide a full evaluation of performance.

An ongoing issue is that evidence on psychometric properties (validity and reliability) of most movement imagery measures is scarce. For imagery generation, self-reported questionnaires are the most used procedure ^10^. Studies usually assess structural validity and reliability of these tools (e.g., factor structure suggesting visual and kinesthetic modalities can be actually generated independently) ^10^, which is in line with consensus-based recommendations ^11^. Some studies also provided evidence on test-retest reliability and measurement error ^12,13^. However, while this should be conducted for all measurement instruments, it is not habitual for other measures of imagery ability, especially behavioural tasks. For maintenance, mental chronometry is commonly used ^14^. In the mental chronometry paradigm, the time required to physically execute an action and the time to imagine it are compared ^15^. Studies investigating the (test-retest) reliability of mental chronometry measures are rare ^16,17^, even though performance is expected to be highly variable over repeated measurements due to learning or fatigue ^18^. Moreover, mental chronometry paradigms are not usually subject of “construct” (internal) validity analyses. That is, whether the paradigms can consistently demonstrate the expected behaviour if movement imagery is being used. For imagery manipulation, a frequent paradigm is the Hand Laterality Judgement Task (HLJT) ^19^, which has been repeatedly used in neuroscientific and clinical research ^20^. In the HLJT, images of hands are presented in different degrees of rotation and in different anatomical views (e.g., palmar or dorsal). To solve the task, the individual may generate an internal representation of their own hand and rotate it to match the on-screen image, which involves an imagined movement (manipulation) of the own hand^21^. The rotation effect shows lower accuracies and longer reaction times for longer compared to shorter rotations^19,22,23^. Further, the ‘biomechanical constraints’ effect (lower reaction times for anatomically possible rotations compared to biomechanically impossible rotations) indicates that movement imagery is being used in the HLJT ^23,24^. However, (test-retest) reliability analyses and measurement error estimation have rarely been conducted in the HLJT ^25^, even though both psychometric properties could be influenced by learning effects due to repeated exposure to experimental stimuli ^26^.

Another significant issue in the field is that studies typically collect data from relatively limited sample sizes ^10^. This substantially contributes to uncertainty regarding the relationship between different measures of imagery (i.e., their criterion or convergent validity). This is typically assessed through correlation analyses under frequentist null-hypothesis significance testing. It has been argued that imagery processes are independent from each other ^6,14,27^, because their outcome measures typically do not *statistically* correlate (*p* > 0.05). Based on our current knowledge, it is therefore possible that an individual self-reports very ‘good’ imagery generation and at the same time performs poorly in maintenance or manipulation tasks ^6^. While this might be naturally plausible if the three processes are truly unrelated and independent, it is a somewhat counterintuitive claim, as we would expect that measures of movement imagery would correlate at least weakly as they all depend upon a ‘common’ skill. The lack of correlation between measures could also be explained by the fact that previous work has been statistically underpowered for these correlation analyses and was unable to detect effect sizes that are actually smaller than expected ^28^. Thanks to technological developments, online studies allow to collect larger, and potentially more representative samples in shorter periods of time than was previously possible ^29^. This has not been generally applied in movement imagery research, even though it might be a potential avenue to address the afore-mentioned limitations ^30,31^. In addition, previous works have used traditional null-hypothesis significance testing, which is not suitable to evidence the *absence* of a correlation. Bayesian statistics would be useful to determine not only the presence of an effect, but also evidence for its absence ^32^.

In this paper we combine three movement imagery tests. We combine measures of generation, maintenance, and manipulation to comprehensively assess movement imagery ability. We leverage online procedures to collect data from a large sample and assess the psychometric properties of the tests, to check validity and reliability with contemporary statistical procedures. Furthermore, we provide the materials through open-source software that requires minimal programming experience to allow the (re-)use of the tests. We hope this will help with standardization of assessment procedures more broadly in the field of movement imagery. Overall, we aim to address fundamental challenges in movement imagery ability assessment by providing psychometrically tested materials.

## METHODS

### Design and ethical compliance

This online study employed a within-subjects, test-retest design. Reporting followed the COSMIN guidelines for measurement instruments ^11^, the GRRAS guidelines for reliability studies ^33^ and the GRASS guidelines for movement imagery research ^34^. Data collection was conducted fully online via the crowdsourcing platform Prolific (https://www.prolific.com/). Ethical approval was obtained by the Ethics Commission of the Institute for Research in the Psychological Sciences at UCLouvain, Belgium (ID: Projet2024-85). All individuals gave explicit electronic informed consent before participating. Participants were financially compensated for their time (10€/session, which lasted _∼_45-75 minutes).

### Participants

Participants were self-reported healthy adults (18-90 years old), with self-reported normal or corrected-to-normal vision and no history of neurological damage. All participants self-reported to be fluent in English, making them able to understand the instructions. Handedness was assessed by an electronic version of the Edinburgh Handedness Inventory ^35,36^. Further, participants reported the hand (left or right) they used to complete the maintenance task.

Participants’ characteristics are summarised in Table 1. A sample of *N* = 180 participants completed the Test session and a subsample of *N* = 123 completed also the Retest, 6.57 ± 0.63 days after (mean ± SD). See Supplementary Fig. S1 for a flowchart showing participants enrolled and excluded after applying the data quality exclusion criteria.

**Table 1.**
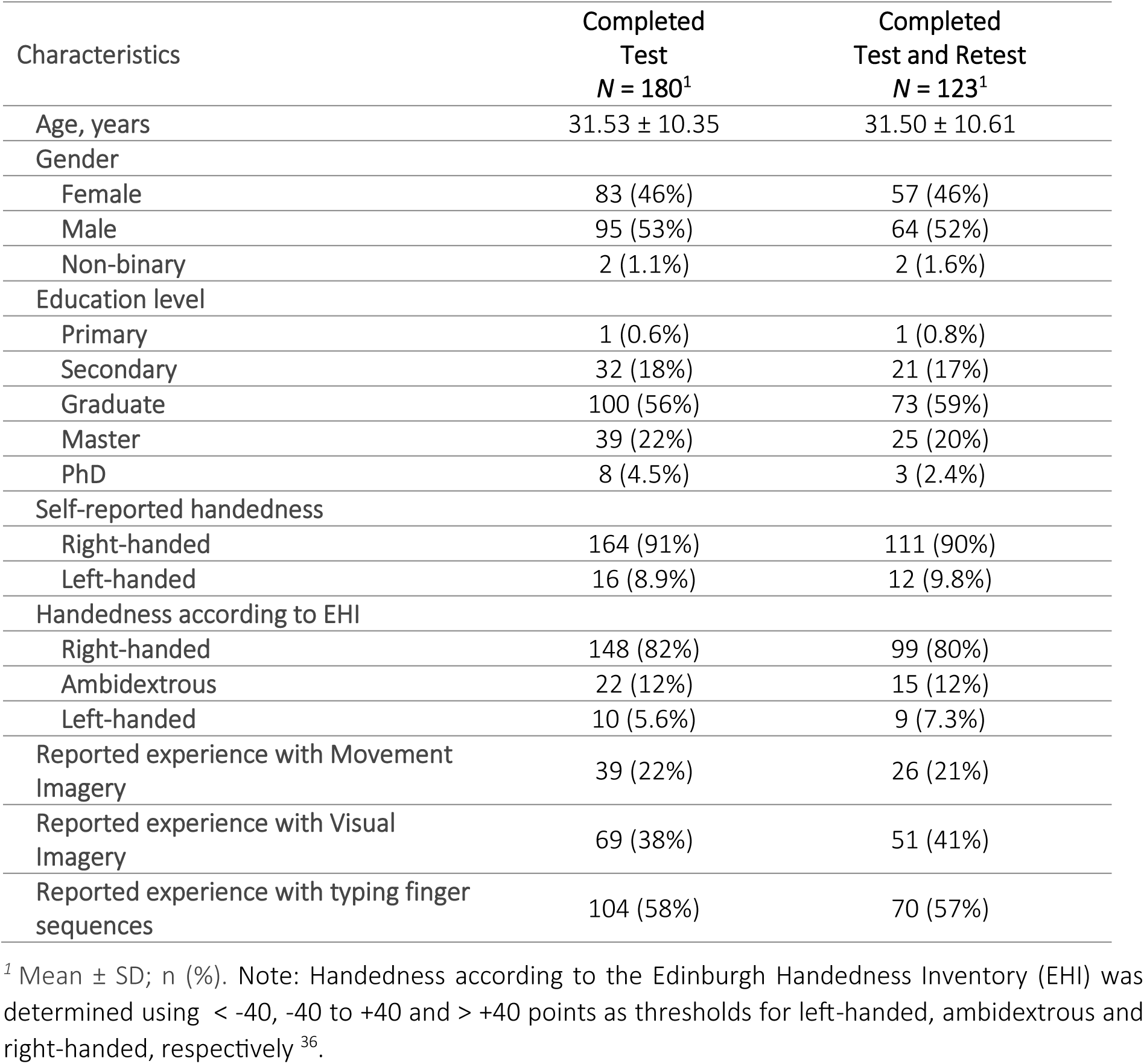
Participants’ sociodemographic characteristics.

### General Experimental Procedure

The task battery (Fig. 1) was created in the open-source software PsychoPy ^37^ version 2024.2.0 (code freely available at https://osf.io/7cfz4/). It was hosted online via Pavlovia (https://pavlovia.org/) ^37^. The electronic questionnaire was created using Pavlovia Surveys and embedded in the PsychoPy experiment. Stimulus size for behavioural tasks was determined using ‘PsychoPy units’, where a size of 1 unit is the height of the screen in landscape orientation.

**Figure 1.**
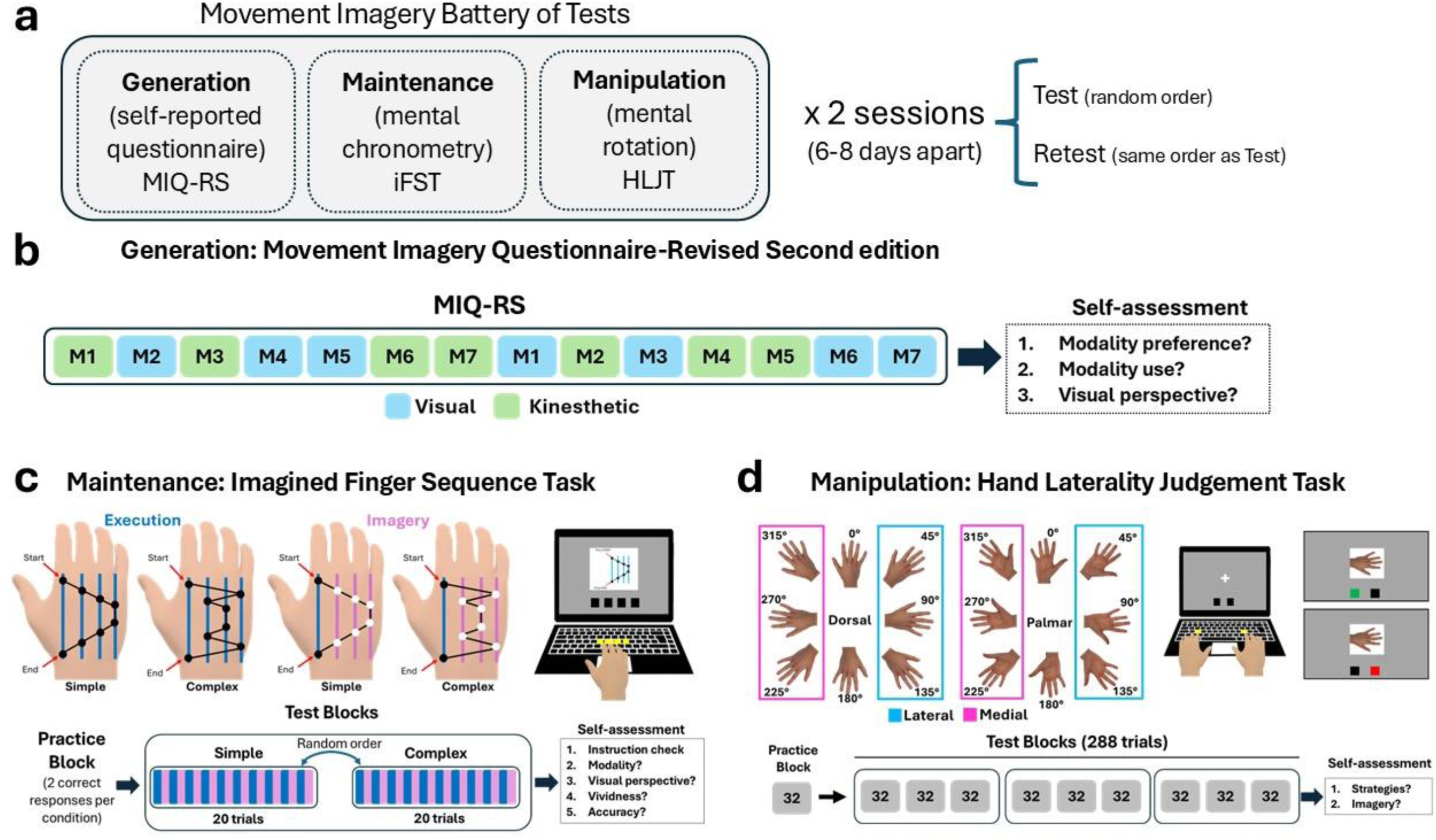
Overview of the tests to assess movement imagery ability. **Panel a** shows the tests to assess movement imagery ability, with measures of generation, maintenance and manipulation. A subset of participants completed the tests twice (6-8 days apart). **Panel b** shows the structure of the MIQ-RS questionnaire to assess generation. Seven movements are imagined in both visual and kinesthetic modalities. Participants then reported their preferences in terms of modalities and visual perspectives, and whether they used the visual/kinesthetic modality when instructed to use kinesthetic/visual only, via single items. **Panel c** shows the structure of the Imagined Finger Sequence Task to assess maintenance. Participants completed two sequences (complex and simple), in two blocks (randomized order) and alternating imagery and execution trials within each block. During imagery, the time was measured by pressing the first element and the last element of the sequence, and the elements in-between were imagined. After each block, participants were asked about the imagery modalities and visual perspectives used, their overall imagery vividness and self-reported accuracy. **Panel d** shows the structure of the Hand Laterality Judgement Task to assess manipulation. Images of rotated hands in 8 possible angles were shown. Participants judged the laterality of the depicted hand (left or right—right-hands are shown in the figure), responding bimanually with their corresponding hand. Stimuli were rotated clockwise or counterclockwise towards medial or lateral orientations. Feedback on accuracy was provided throughout the task via two small boxes. After the task, participants described their strategies with an open question and were asked about their use of imagery (sensory modalities and visual perspectives).

Participants who completed the whole experiment twice were instructed to maintain approximately the same conditions between Test and Retest (e.g. level of tiredness and attention, which were asked in both sessions on 11-point rating scales—see Supplementary Materials), as well as using the same computer, and to be in a quiet place.

Each session, participants were evaluated by three types of assessments of movement imagery ability: a questionnaire to assess generation, a finger sequence task to assess maintenance, and the HLJT to assess manipulation. The order of the assessments was random for each participant, though participants who completed both the Test and Retest did it in the same order.

### Generation—Questionnaire

The Movement Imagery Questionnaire-Revised Second Edition (MIQ-RS) was used ^13^, as it was developed for people with motor impairments and thus would enhance clinical applicability of the resource in the long term. The MIQ-RS is a 14-item self-administered tool that assesses 7 movements (hip and knee flexion, shoulder flexion, horizontal shoulder adduction, trunk flexion, pushing a swinging door, grasp a glass and opening a door) in 2 sensory modalities: visual imagery and kinesthetic imagery. Note that for visual imagery it was not specified whether the first- or third-person perspective should be used. Each item involves 4 steps: (1) adopting an initial position; (2) physically performing a movement one time; (3) returning to the starting position; and (4) visually or kinesthetically imaging the movement one time. After that, participants rate the ease of generating the image on a 7-point Likert scale from 1 = very hard to see/feel to 7 = very easy to see/feel. Visual and kinesthetic items are interspersed (see Fig. 1b). To minimise errors when switching between modalities, we color-coded each item instruction according to the intended modality. An attention check question was interleaved with actual items approximately half-way through, allowing us to exclude participants who missed it. As individual-level indicators of performance, we obtained visual and kinesthetic subscale sum-scores (7-49 points), where higher scores indicate better ability.

#### Self-assessments

After completing the MIQ-RS, participants were asked about their sensory modality preferences (visual/kinesthetic), if they involuntarily switched between modalities, and their preference for visual perspective(s) during the visual items, on 11-point scales (see Supplementary Materials).

### Maintenance—Mental chronometry task

We developed a new behavioural paradigm, which we named the Imagined Finger Sequence Task (iFST). The task was developed to overcome limitations of previous mental chronometry paradigms, in which no clear manipulation checks can be conducted for internal validity analyses ^38^.

The task consisted in typing, or imagining typing, two 8-digit sequences with the index, middle, ring and little fingers of the dominant hand (digits 2, 3, 4 and 5). The ‘F’, ‘G’, ‘H’ and ‘J’ keys were used as they are centred in the keyboard and their location is consistent across different montages (AZERTY/QWERTY). For right-handed participants, the F, G, H and J keys corresponded to the index, middle, ring and little finger, and the opposite for left-handed. To minimise the use of explicit cognitive strategies (e.g., ‘verbal repetition’) rather than imagery in sequence recall, no explicit mention to a key-digit or digit-number association was offered. Two types of sequences were employed based on their complexity, always with 2 repetitions per digit. The ‘simple’ sequence used a staircase design, with 1 change in direction: 2-3-4-5-5-4-3-2 (see Fig. 1c). The ‘complex’ sequence was designed to be more challenging, including 4 changes in direction but the same number of repetitions per digit: 2-5-3-4-4-3-5-2. We expected the duration of execution and imagery to be longer for the complex than the simple sequence, which was the ‘manipulation check’ in this task. Additionally, it allowed us to construct individual-level measures overcoming traditional limitations of using absolute deviations from execution as an outcome measure in mental chronometry ^38^. Both sequences started and finished with the index finger to allow comparison with the imagery condition—see below. The sequences were presented via diagrams (Fig. 1c). Before the main task blocks, participants had a familiarization block of the execution and imagery conditions for each sequence, where they received online visual feedback on key presses through 4 small boxes placed at the bottom of the screen (0.07 x 0.07 PsychoPy units) and offline trial-to-trial feedback on response accuracy. To advance from familiarization to the test blocks, participants needed at least 2 correct responses per condition.

During test blocks, participants were asked to either physically execute the sequence or to imagine executing the sequence. They were asked to (imagine to) perform the sequence as quickly and accurately as possible, but prioritising accuracy over speed. In the imagery condition, participants were instructed to actually press the first key and the last key with the index finger. They imagined all the key presses in-between without physically executing any finger movements. They were instructed to synchronize the first and last executed key presses with their mental simulation (i.e., as if the key was the first/last element of the action). Participants were asked to perform imagery combining a first-person visual modality (i.e., as if they were seeing the movement through their own eyes) and a kinesthetic modality (i.e., focusing on the feeling of the movement). Sequences (simple and complex) were performed in blocks (counterbalanced order). Each block, participants performed 10 execution and 10 imagery trials of a given sequence type. Each imagery trial was preceded by an execution trial. Therefore, participants performed 20 trials per sequence (10 execution and 10 imagery), for a total of 40 trials. During the test blocks, no feedback was provided.

As indicators of performance, two traditional mental chronometry scores were computed for each individual, where higher scores indicate lower imagery ability ^39^. The absolute raw difference (in seconds) between execution and imagery:

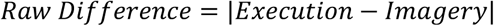

The absolute relative difference (% of estimation error from execution):

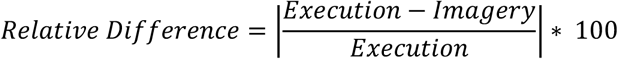

Both scores were computed for simple and complex sequences separately and then averaged to obtain a ‘global’ performance score per individual. As deviations from real-time movement imagery in mental chronometry tasks may be difficult to interpret ^38^, we also obtained a more principled score, known as the constraint approach ^7^. In this framework, movement imagery ability is approximated via the proportional change in execution and imagery durations after introducing a constraint to the action. These proportional changes in execution and imagery durations are subtracted, such that a smaller difference indicates higher movement imagery ability. In our procedure, the constraint is the use of a complex sequence compared to the simple sequence. This indicates how well imagery retains the constraint compared to the constraint’s influence on physical execution. Therefore, the formula takes the time needed to complete complex and simple sequences for execution (Exe) and imagery (Ima):

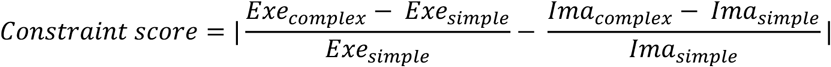

#### Self-assessments

After each block, participants were asked which imagery modality they were indicated to use (as an instruction check question), their modality preference, visual perspective, vividness, and perceived accuracy during imagery on 11-point scales (see Supplementary Materials).

### Manipulation—Hand Laterality Judgement Task (HLJT)

The paradigm was the same as a recently developed version by our lab ^31,40^. Participants decided whether a picture of a hand corresponded to the left or right side. Experimental stimuli were real hand images (Fig. 1d) presented in 8 rotational angles in the frontal axis (0°, 45°, 90°, 135°, 180°, 225°, 270°, 315° for right hands in the clockwise direction, and the opposite for left hands), and 2 possible views (palmar or dorsal). In total, 32 unique stimuli were used (8 angles x 2 views x 2 hand sides). Biomechanical constraints were considered as the difference between medial and lateral rotations (Fig. 1d), indicative of motor processing in this task ^22,23,41^.

Each stimulus was preceded by a fixation cross in the centre of the screen for 800ms. Stimuli were presented in the centre and remained until a response was made. Participants responded bimanually using the index finger of their left or right hand, mapping left/right responses to left/right images (Fig. 1d). Trial-to-trial visual feedback was provided for 300ms via two small boxes at the bottom of the screen, which turned green/red if a response was correct/incorrect. Participants were allowed to familiarize with task and stimuli in a familiarization block with 32 trials (1 repetition per unique stimulus). Then they completed 3 test blocks and 96 trials per block. In each test block, stimuli were randomly presented in sub-blocks of 32 trials with 1 repetition per unique stimulus in each sub-block, to minimize the likelihood of the same stimulus appearing consecutively. The 3 test blocks resulted in a total of 288 trials (9 repetitions per unique stimulus). Breaks between blocks lasted at least 10 seconds. As indicators of performance, the outcome measures were the overall accuracy (% of correct responses) and reaction time (ms).

#### Self-assessments

After completing the HLJT, participants were qualitatively debriefed about their strategies (open question), if they used movement imagery in the task explicitly (yes/no), and which modalities and perspectives of imagery they used. It is important to mention that because the order in which participants completed the tasks was random, 1/3 of participants completed the HLJT as the first test. We wanted to know whether participants would report using imagery in the HLJT without being explicitly told that the study was about movement imagery. The third of participants who completed the HLJT first, were not aware that the study was about imagery (all participants were told the study was about “measuring cognitive abilities”, without explicitly mentioning imagery until they completed the MIQ-RS or the iFST). This allowed us to compare the responses to these self-assessments of the HLJT between those participants aware of the concept of imagery before completing the task, and those unaware. See Supplementary Materials for details.

### Data pre-processing and Exclusion criteria

A flow diagram illustrating this process is shown in Fig. S1.

#### MIQ-RS

Item-level responses were collected, and participants were excluded if they responded incorrectly to the attention check question.

#### iFST

The execution time and imagery time (time elapsed between the first and last key presses, in seconds), as well as response accuracy, were collected. For execution, only trials with correct complete sequences (all 8 key presses) were considered. For imagery, only trials with correct responses (2 key presses with the indicated start and finish keys) were considered. Participants were excluded if they: 1) had <75% overall correct responses, for both execution and imagery separately, meaning they did not prioritise accuracy over speed in the task; or 2) the average execution or imagery time was <1s, reflecting they did not appropriately perform the task. We performed sensitivity analyses excluding participants who responded incorrectly to any of the 2 instruction checks but did not use this as an exclusion criterion (see Supplementary Materials).

#### HLJT

Accuracy (% of correct responses) and reaction time were analysed separately. First, trials where reaction time was <300MS or <3,000ms, likely reflecting anticipatory responses and no engagement with the task, respectively, were discarded ^31^. For reaction time, only trials with correct responses were considered. We excluded all participants who had: 1) <60% overall accuracy, indicating their performance was not above chance level (50%); or 2) >50% trials rejected due to short or long reaction times.

### Psychometric Assessment & Statistical Analysis

Analyses were conducted in R version 4.4.3 (R Core Team 2025). Data and scripts are freely available at https://osf.io/7cfz4/. For statistical significance, the Type I error rate was fixed at 5%. For parameter uncertainty, 95% confidence intervals (95%CI) were obtained, unless indicated otherwise.

#### Structural validity and reliability (MIQ-RS)

The internal structure was first assessed by Confirmatory Factor Analysis via maximum likelihood estimation (package ‘lavaan’ v0.6-19 ^42^). Model fit indices included the χ² test for the baseline model, Comparative Fit Index (CFI), Tucker-Lewis Index (TLI), Root Mean Square Error Approximation (RMSEA) with a 90%CI and Standardized Root Mean Square Residual (SRMR). First, a model with the original two-factor structure was tested. However, the model showed poor fit (see Results) due to residual correlations between visual and kinesthetic items representing the same movement (e.g., item 1 and item 8—see Fig. 1b). Given that ratings for visual and kinesthetic imagery of the same movement would be expected to covary (representing action-specific covariance) ^43^, we specified residual correlations between items that represent the same movement across kinesthetic and visual modalities. The final model showed substantially better fit (see Results). As the factorial structure of the questionnaire was confirmed, the next analyses were conducted for both subscale scores independently. Reliability was tested as internal consistency, using hierarchical Omega (ω) coefficient, which is optimal when data are non-normally distributed, and the questionnaire is not unifactorial or tau-equivalent ^44^. 95%CIs were calculated with bias-corrected and accelerated bootstrap (500 replicates) using package ‘MBESS’ v4.8.1. A value of ω > 0.7 was considered acceptable ^44^.

#### Internal validity of behavioural tasks (iFST and HLJT)

First, we tested for naturally expected effects of both paradigms at the group level that would indicate embodied cognition. Second, we performed psychometric testing at the individual level. Generalised linear mixed-effects models (GLMMs) with full random effect structure (intercepts and slopes for participants) were used, alongside Bonferroni-corrected post-hoc tests. Predictions were reported back-transformed from the link scale as appropriate (e.g. from log-link to log scale) ^45^.

- For the iFST, we first checked for differences in accuracy (% correct) between simple and complex sequences for execution trials via a binomial GLMM. For the main analysis, we did expect an effect of sequence complexity on both execution and imagery times. Therefore, we assessed whether the time to complete the trial was different between Sequences (simple vs complex) for the different Types of trials (execution and imagery). A GLMM with a gamma distribution was used, because time was right-skewed. Predefined post-hoc tests assessed differences between sequences for each type of trial. Finally, we also assessed learning effects in this task (see Supplementary Materials).
- For the HLJT, we expected a strong effect of rotation Angle for both accuracy and reaction time, and an interaction with hand View (palmar vs dorsal) ^31,40^. To confirm these effects, two GLMMs, were used with binomial (accuracy as binary 0-1 for each trial) and gamma (reaction time in ms, right-skewed) distributions. Predefined post-hoc comparisons tested differences between sequential pairs of angles within each view (0° vs 45°, 45° vs 90°, etc), and medial vs lateral pairs (45° vs 315°, 90° vs 270°, 135° vs 225°) testing the biomechanical constraints effect for each view. We predicted differences would be greater in the palmar view than the dorsal view ^31,40^. Additionally, we expected the presence of the biomechanical constraints effect for reaction time overall, in the palmar view, but not in the dorsal view. For this we ran a separate GLMM including trials with medial and lateral rotations only (i.e., without 0° and 180°), with post-hoc tests between each level of direction (medial vs lateral) for each level of view (palmar and dorsal).

#### Reliability of behavioural tasks (iFST and HLJT)

We used generative Bayesian hierarchical models to estimate the reliability (signal-to-noise ratio—SNR) of the experimental conditions in each task ^46^. We used full random effects hierarchical models mimicking our frequentist GLMMs above (using the same formulas). For the iFST, the model predicted the time to complete sequence, with sequence complexity and trial type as predictors. For the HLJT, the model tested the biomechanical constraints effect on reaction time with direction (medial vs lateral) and hand view (dorsal vs palmar) as predictors. The models were run with uninformative (but not uniform) priors, using ‘brms’ package v2.22. Full details are explained in the Supplementary Materials. We extracted SNRs as the ratio between the variability of participants’ average effects (signal) and their average trial-level variability (noise) for each parameter ^47^. SNR > 1 and SNR < 1 indicated higher and lower reliability, respectively.

#### Criterion validity

Spearman’s correlation coefficients were used to assess the relationship between the three processes assessed by the tests. We used Spearman’s correlation coefficients as the majority of the measures did not show normal distributions. To overcome previous limitations of frequentist analyses to provide evidence *for the absence* of an effect, we used Bayesian correlations. Coefficients were interpreted as negligible, weak, moderate, strong or very strong correlation if *r* was <0.1, 0.1-0.4, 0.4-0.7, 0.7-0.9 and >0.9, respectively ^48^. Uninformative (wide) priors were used (beta 1.73 ± 1.73), and Bayes Factors (BFs) are reported in favour of the null (*BF_01_*) or alternative (*BF_10_*) hypothesis, with established benchmarks for interpretation ^49,50^ (package ‘correlation’ v0.8.8). The individual-level outcome measures were MIQ-RS subscale sum-scores (visual and kinesthetic), the three scores of the iFST (raw difference, relative difference and constraint score), and the two scores of the HLJT (accuracy, reaction time). Correlations *between* tests, but not between scores *within* a test were evaluated.

#### Test-retest reliability and measurement error

For all outcome measures, test-retest reliability was calculated by the Intraclass Correlation Coefficient (ICC), using a two-way mixed-effects model with absolute agreement and single measures (2,1) ^51^. Acceptable values were ICC > 0.5, and ICCs were interpreted as poor, moderate, good and excellent reliability if ICC < 0.5, 0.5-0.75, 0.75-0.9 and > 0.9, respectively ^51^. Limits of agreement were assessed with differences between Test and Retest plotted against the means of the two measurements by Bland-Altman plots ^52^. Additionally, measurement error was tested using standard error of measurement (SEM = SD * √(1 – ICC)) and minimal detectable change with 95% confidence level (MDC_95_ = SEM * 1.96 * √2) ^53,54^. Sensitivity analyses were performed for all these measures, considering the level of tiredness/fatigue and attention of the participant from Test to Retest (see Supplementary Materials).

### Sample Size Calculations

Sample size estimations were based on a power of 80% and 5% Type I error rate in a frequentist framework. Calculations are explained in detail in Supplementary Materials. Briefly, the strictest estimation (confirmatory factor analysis of MIQ-RS) required *N* = 176 participants. Therefore, N=180 participants for the Test session were recruited, as it would give adequate power for all our pre-planned analyses.

A separate precision-based sample size calculation was performed for the test-retest reliability. Considering 2 measurements (Test and Retest), an expected ICC = 0.7 (moderate reliability) and a 95%CI width of 0.2 (ICC ± 0.1), *N* = 105 participants were required (package ‘presize’ v0.3.7 ^55^). Note that the expected ICC is conservative for the MIQ-RS ^13,56^ or the HLJT ^25^ but expected for the iFST outcomes, from which we had no prior data.

## RESULTS

At Test, participants took 51.79 ± 14.64 minutes to complete the whole experiment (MIQ-RS: 11.85 ± 7.31 minutes; iFST: 17.38 ± 7.32 minutes; HLJT: 22.55 ± 8.42 minutes). Descriptive statistics for the tests on the first session are shown in Table 2.

**Table 2.** Descriptive statistics for the different tests (*N* = 180).

| Test | Variable | Mean | SD | Min | Max | Median | Q1 | Q3 | IQR |
| --- | --- | --- | --- | --- | --- | --- | --- | --- | --- |
| MIQ-RS | Visual subscale (score) | 38.34 | 6.79 | 7.00 | 49.00 | 39.00 | 35.00 | 43.00 | 8.00 |
|  | Kinesthetic subscale (score) | 34.35 | 8.02 | 9.00 | 49.00 | 35.00 | 30.75 | 39.25 | 8.50 |
| iFST | Simple sequence, execution (s) | 3.05 | 1.14 | 0.90 | 8.30 | 2.88 | 2.26 | 3.58 | 1.32 |
|  | Simple sequence, imagery (s) | 2.97 | 1.27 | 0.23 | 8.29 | 2.78 | 2.18 | 3.64 | 1.45 |
|  | Complex sequence, execution (s) | 4.20 | 1.93 | 1.46 | 11.50 | 3.71 | 2.87 | 4.90 | 2.03 |
|  | Complex sequence, imagery (s) | 3.47 | 1.60 | 0.46 | 13.41 | 3.21 | 2.50 | 4.16 | 1.66 |
| HLJT | Accuracy (%) | 0.90 | 0.08 | 0.61 | 1.00 | 0.92 | 0.88 | 0.96 | 0.08 |
|  | Reaction time (ms) | 1207.49 | 316.36 | 573.45 | 2371.17 | 1163.50 | 980.39 | 1380.80 | 400.41 |
|  | Biomechanical constraints, dorsal (ms) | -53.54 | 153.72 | -486.86 | 454.77 | -36.79 | -156.89 | 46.23 | 203.12 |
|  | Biomechanical constraints, palmar (ms) | -214.99 | 171.89 | -671.79 | 216.94 | -190.22 | -323.15 | -96.49 | 226.67 |

*Note:* In iFST and HLJT, data comes from collapsing across trials/conditions. The biomechanical constraints are calculated as the difference between the mean reaction time in medial orientations minus lateral orientations.
Abbreviations: HLJT: Hand Laterality Judgement Task; iFST: Imagined Finger Sequence Task; IQR: Interquartile Range; MIQ-RS: Movement Imagery Questionnaire-Revised Second edition; Q1/Q3: First/Third quartiles.

### Structural validity and reliability of the MIQ-RS

The original factor analysis model had a poor fit (χ²(76) = 229.03, *p* < 0.001; CFI = 0.85; TLI = 0.82; RMSEA = 0.11 90% CI [0.09, 0.12]; SRMR = 0.07). After allowing for residual correlations between items with the same movement across visual and kinesthetic modalities, the new model had adequate fit indices (χ²(67) = 112.57, p < 0.001; CFI = 0.955; TLI = 0.939; RMSEA = 0.061 90% CI [0.041, 0.081]; SRMR = 0.057). In the movement-specific model, the two factors representing visual and kinesthetic subscales were moderately correlated (*r* = 0.43), and factor loadings varied from λ = 0.59-0.7 for visual items (except item 5: λ = 0.48) and λ = 0.66-0.78 for kinesthetic items (Fig. 2a). Reliability (internal consistency) was adequate (ω = 0.83 95%CI [0.73 0.89]). Item-total correlations and correlations after removing items did not show issues with any specific items (all *r* > 0.3; Table S1). The distributions of subscale sum-scores are shown in Fig. S2, showing slightly higher group-level scores for visual imagery (*mean* ± *SD* = 38.34 ± 6.79) than for kinesthetic imagery (34.35 ± 8.02).

**Figure 2.**
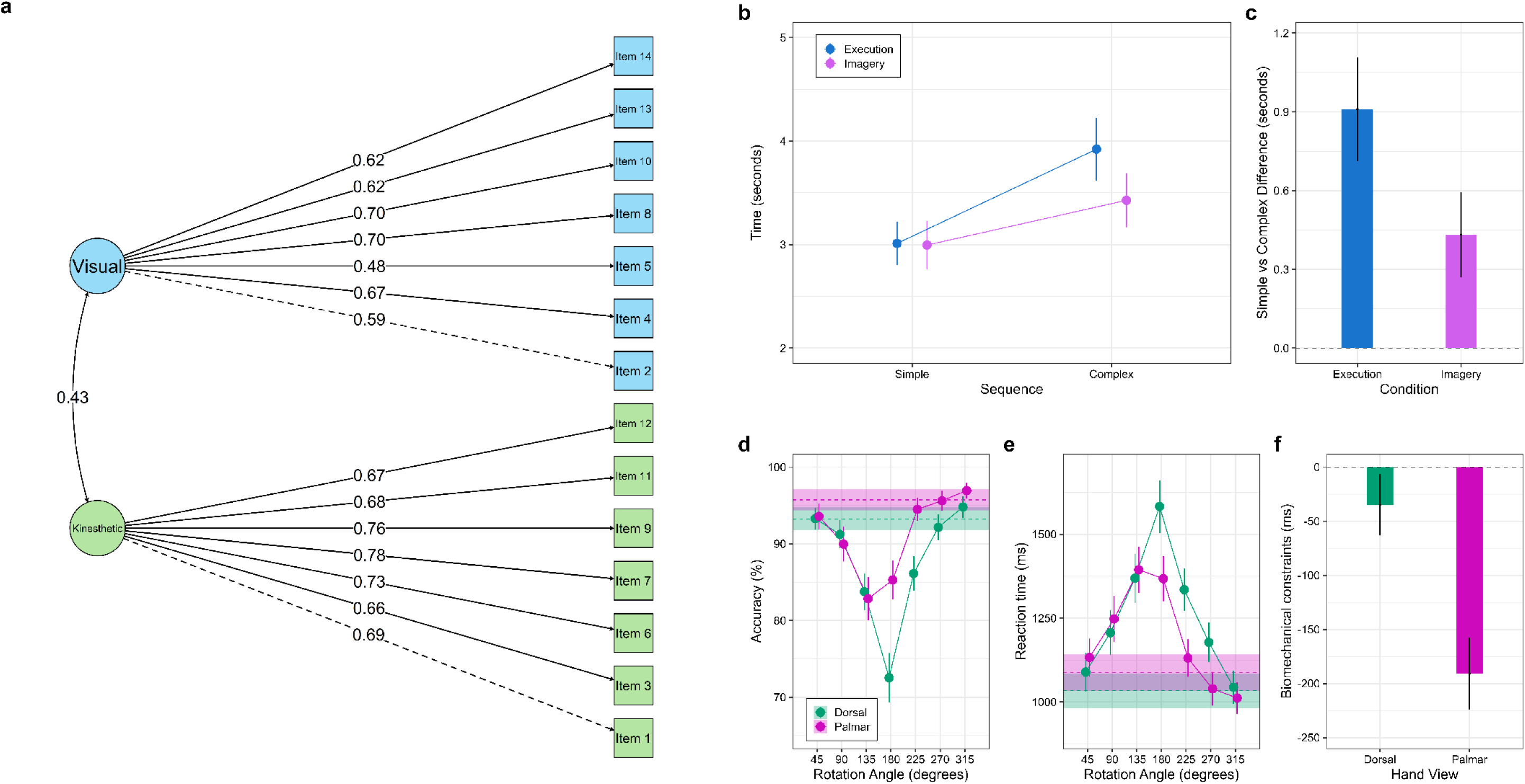
Main results of the structural and internal validity analyses. **Panel a)** shows the results of the Confirmatory Factor Analysis of the Movement Imagery Questionnaire-Revised Second edition. Kinesthetic and visual items loaded onto their corresponding latent variables (kinesthetic and visual subscales, which showed a moderate factor-factor correlation). Factor loadings ranged from 0.48 to 0.7 for the visual subscale and from 0.69 to 0.78 for the kinesthetic subscale. **Panels b)** and **c)** show the results of the generalised linear mixed model of the Imagined Finger Sequence Task. **Panel b)** shows model-estimated marginal means for each trial type (execution and imagery) and for each sequence type (simple and complex). For both trial types, the complex sequence required more time to complete than the simple sequence, as predicted. However, the constraint of the complex sequence was stronger for execution than imagery, as shown in **Panel c)**. **Panels d-f** show the results of the generalised linear mixed models of the Hand Laterality Judgement Task. **Panels d)** and **e)** show model-estimated marginal means for each rotation angle compared to the baseline (horizontal dashed line shows data for 0° degrees of rotation and its 95%CI) for each hand view. A clear symmetric shape around 180° was found for the dorsal view in both accuracy and reaction time. A clear asymmetric shape was found for the palmar view in both measures, indicating the presence of a biomechanical constraints effect primarily in this view. This was confirmed by the separate model as shown in **Panel f)**, which illustrates the ‘grand’ biomechanical constraints effect (medial vs lateral difference) in palmar and dorsal views separately.

### Internal validity and reliability in the iFST

Accuracy on the iFST was on average 95.2% [94.6, 95.8], consistent with the instructions to prioritize accuracy over speed. The binomial GLMM did not provide evidence for a difference in accuracy for execution trials between simple sequences (92.19% [90.27, 94.11]) and complex sequences (93.99% [92.41, 95.58]), difference = -1.8% [-4.2, 0.06], *z* = -1.48, *p* = 0.1402.

For time to complete the sequence, the GLMM included 6,679 trials after data trimming, which removed 2.57% of trials. For execution (Fig. 2b), the model estimated an average of 3.01 seconds [2.8, 3.22] to complete the simple sequence and 3.92 seconds [3.62, 4.22] to complete the complex sequence, with evidence for a difference (Fig. 2c) between the two (post-hoc mean difference = -0.91 seconds [-1.11, -0.71], *t*_(6664)_ = -9.02, *p* = 4.68×10^-19^). For imagery, the average time to complete the simple sequence was 3.00 seconds [2.76, 3.23] and for the complex sequence 3.43 seconds [3.16, 3.69], with evidence for a difference but smaller (post-hoc mean difference = -0.43 seconds [-0.59, -0.27], *t*_(6664)_ = -5.21, *p* = 3.88×10^-7^). These results demonstrate the internal validity of the paradigm.

For reliability, data regarding the SNRs from the hierarchical Bayesian model are shown in Table 3 (and Fig. S3) and showed adequate to moderate reliability.

**Table 3.** Signal-to-noise ratios (SNRs) estimated from the generative hierarchical Bayesian models. For each parameter in each task, the SNR was calculated as the Signal (average inter-individual variability for an effect across participants) and Noise (average intra-individual variability for an effect across trials) from the posterior distribution.

| Imagined Finger Sequence Task (iFST) |  | Hand Laterality Judgement Task (HLJT) |  |
| --- | --- | --- | --- |
| Parameter | SNR | Parameter | SNR |
| Intercept | 8.82 | Intercept | 11.6 |
| Sequence | 2.82 | Direction | 1.75 |
| Trial type | 4.92 | Hand view | 2.82 |
| Sequence x Trial type | 1.58 | Direction x Hand view | 1.76 |

### Internal validity and reliability in the HLJT

The accuracy GLMM included 49,255 trials after data trimming (4.98% removed trials). The reaction time GLMM included 44,468 correct trials (90.2% accuracy overall). The models showed a significant effect of rotation Angle and interaction with hand View.

For accuracy, the effect of rotation was non-monotonic (Fig. 2d), as differences between sequential pairs of angles (0° vs 45°, 45° vs 90°, etc.) varied in magnitude depending on the comparison pair (Table S2), in both palmar and dorsal views. The interaction Angle by View showed that lateral rotations (45°, 90°, 135°), were processed significantly less accurately than their medial counterparts (315°, 270°, 225°) in the palmar view (45° vs 315°: difference = 3.35% [1.66, 5.05], *z* = 3.88, *p* = 1.05×10^-4^; 90° vs 270°: difference = 5.66% [3.41, 7.91], *z* = 4.93, *p* = 8.27×10^-7^; 135° vs 225°: difference = 11.65% [8.83, 14.47], *z* = 8.09, *p* = 5.95×10^-16^), but not in the dorsal view (differences ≤ 2.38%, *z* ≤ 1.81, *p* ≥ 0.07). This illustrated a biomechanical constraints effect in the palmar view.

For reaction time, the results were similar in terms of a non-monotonic effect of rotation angle (Fig. 2e, Table S3). Lateral rotations were processed significantly faster than medial directions, specifically in the palmar view (45° vs 315°: difference = -120.79ms [-150.56, -91.03], *t*_(44315)_ = - 7.95, *p* = 1.85×10^-15^; 90° vs 270°: difference = -208.25ms [-255.46, -161.04], *t_(44315)_* = -8.64, *p* = 5.5×10^-18^; 135° vs 225°: difference = -263.25ms [-309.55, -216.95], *t*_(44315)_ = -11.14, *p* = 8.32×10^-29^). In the dorsal view significant differences between directions were only found comparing 45° vs 315° (difference = -45.63ms [-76.39, -14.88], *t*_(44315)_ = -2.91, *p* = 0.003), but not 90° vs 270° (difference = -28.37ms [-67,91, 11.16], *t_(44315)_* = -1.4, *p* = 0.159) or 135° vs 225° (difference = -34.38ms [-74.16, 5.39], *t*_(44315)_ = -1.69, *p* = 0.09).

The biomechanical constraints GLMM included 33,885 correct trials (95.64% trials after excluding 0° and 180°). The model showed a strong interaction between Direction and View (Fig. 2f). Overall, medial directions showed shorter reaction times than lateral directions in the palmar view (medial = 1060.29ms [1010.82, 1109.76]; lateral = 1251.17ms [1191.54, 1310.8] and the dorsal view (medial = 1179.84ms [1125.7, 1233.98]; lateral = 1214.69ms [1151.68, 1277.70]. The difference illustrating the ‘grand’ biomechanical constraints was significant in both views, but larger in the palmar view (difference = -190.88ms [-223.98, -157.77], *t_(33870)_* = - 11.3, *p* = 1.45×10^-29^) than in the dorsal view (difference = -38.84ms [-63.1, -8.59], *t_(33870)_* = -2.42, *p* = 0.016).

In terms of reliability, the hierarchical Bayesian model (Table 3, Fig. S4) showed moderate reliability estimates.

### Criterion validity

The MIQ-RS visual subscale showed a weak correlation with the raw difference in the iFST (*r* = -0.15 [-0.29, -0.1]), but evidence was inconclusive (*BF_10_* = 1.07). The kinesthetic subscale weakly correlated with the accuracy in the HLJT (*r* = 0.17 [0.03, 0.31]) with moderate evidence against the null (*BF_10_* = 1.72). The raw difference in the iFST weakly correlated with the reaction time in the HLJT (*r* = 0.27 [0.13, 0.4]) with extreme evidence against the null (*BF_10_* = 141.32) and the relative difference correlated with a weaker effect size (*r* = 0.18 [0.04, 0.33]) and moderate evidence (*BF_10_* = 2.34). For the rest of the correlations, evidence was generally moderate in favour of the null hypothesis of no correlation (*BF_01_* > 3) or inconclusive results (*BF_01_* ≈ 1). Overall, the absence of correlations between tests generally showing weak to negligible effect sizes (Fig. 3, Table S4), indicated that the tests measure distinct aspects of movement imagery ability.

**Figure 3.**
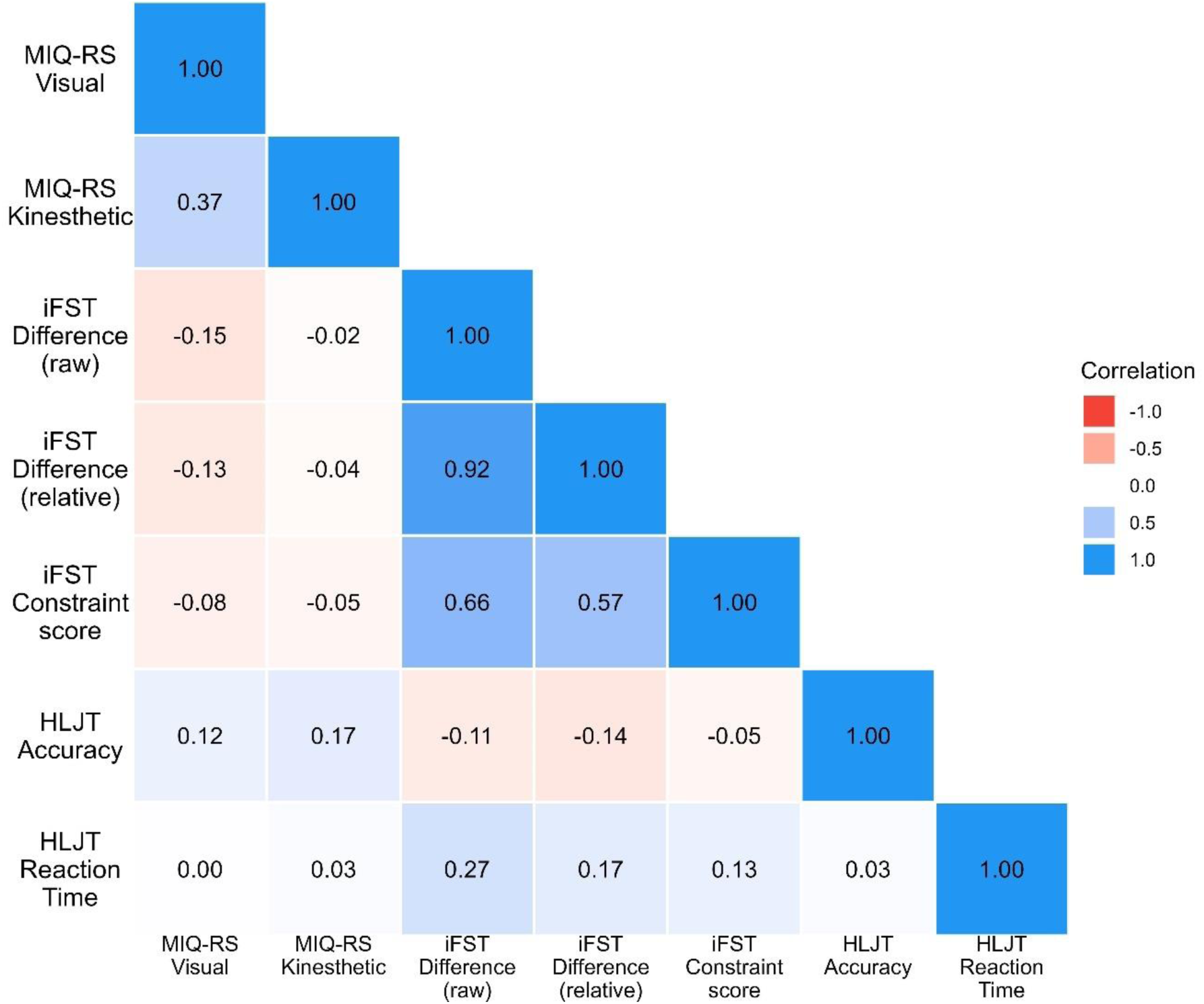
Heatmap showing Bayesian Spearman’s rank correlations between the three tests of the movement imagery ability assessment. Generation was assessed as visual and kinesthetic subscale sum-scores of the Movement Imagery Questionnaire-Revised Second edition (MIQ-RS). Higher scores indicated better imagery ability. Maintenance was assessed as the raw and relative difference between execution and imagery or the difference in inducing a constraint to the action between them, by the Imagined Finger Sequence Task (iFST). Higher scores indicated poorer imagery ability (greater differences between imagery and execution). Manipulation was assessed as the average accuracy and reaction time in the Hand Laterality Judgement Task (HLJT). Higher values of accuracy, and lower values of reaction time indicated better imagery ability.

### Test-retest reliability and measurement error

Results are shown in Table 4 and Figs. S5-S7. Overall, ICCs showed moderate to good test-retest reliability (*ICCs* ≥ .67, except for the constraint score). For the MIQ-RS, the kinesthetic subscale showed higher ICC compared to the visual subscale (*ICC_(2,1)_* = 0.81 [0.75, 0.87]; *ICC_(2,1)_* = 0.67 [0.56, 0.76]). For the iFST, the two mental chronometry scores had similar ICCs (*ICC_(2,1)_* = 0.81 [0.72, 0.87]; *ICC_(2,1)_* = 0.77 [0.69, 0.84]), but the Constraint score showed unacceptable reliability (*ICC_(2,1)_* = 0.33 [0.16, 0.48]). For the HLJT, both accuracy and reaction time showed good reliability (*ICC_(2,1)_* = 0.78 [0.63, 0.87]; *ICC_(2,1)_* = 0.79 [0.34, 0.91]) as well as the biomechanical constraints effect (*ICC_(2,1)_* = 0.69 [0.59, 0.78]). These results were qualitatively equivalent (Table S5) after excluding participants reporting differences between Test and Retest in terms of their level of tiredness/fatigue and/or attention (difference >2 points on 0-10 rating scales), a sensitivity analysis which excluded *N*=22 participants.

**Table 4.**
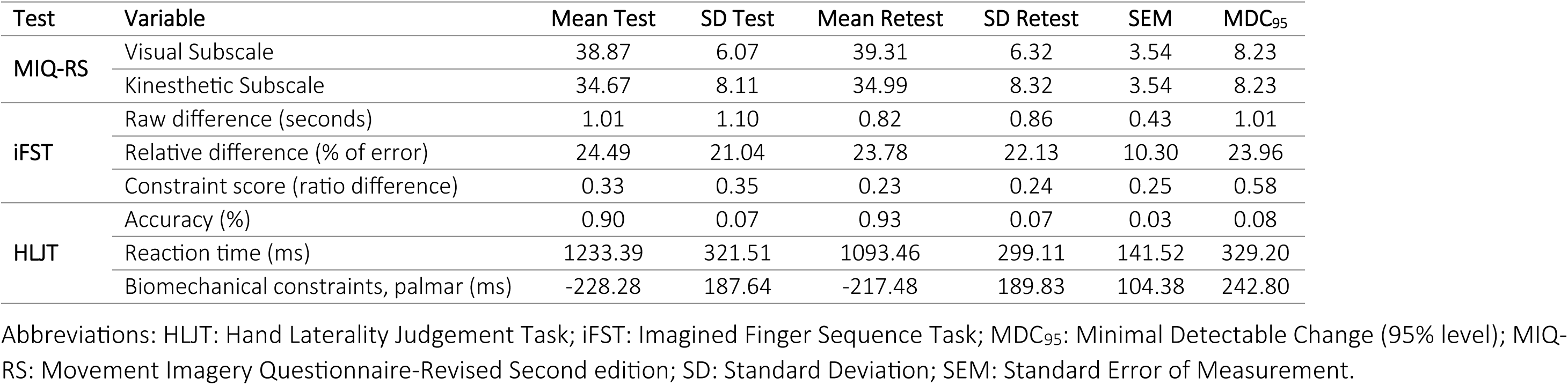
Change from Test to Retest in performance and measurement error results (*N* = 123).

### Self-assessments

The MIQ-RS results revealed that participants preferred a visual modality or had no preference over kinesthetic and similarly favoured a first-person visual perspective over the third-person (Fig. S8). When performing visual items, kinesthetic imagery was used minimally or moderately, whereas visual imagery was used moderately to highly during kinesthetic items. In the iFST, participants reported moderate to high vividness across sequences, with slightly lower perceived accuracy for complex sequences but overall, few self-reported errors (Fig. S9). Use of both visual and kinesthetic modalities was moderate to high, with a strong preference for first-person perspective and very low use of third-person perspective. In the HLJT, most participants (>70%) reported thinking about their hands and using imagery regardless of prior instruction or awareness about the concept of imagery (Fig. S10). Visual modality use was high to very high, kinesthetic use varied (bimodal distribution), and first-person perspective was strongly preferred over third-person (Fig. S11).

## DISCUSSION

This study investigated the psychometric properties of three tests of movement imagery ability focusing on generation, manipulation and maintenance. The study collected the data fully online. Globally, the results showed adequate validity and reliability of the different tests. Importantly, our results are consistent with the claim that the different tests evaluate separable processes of imagery ^6^, as shown by the general absence of correlations between their scores.

### MIQ-RS

Our findings support the structural validity and reliability of the MIQ-RS in an online sample, extending previous laboratory-based studies ^13,56,57^. The improved model fit after accounting for residual correlations between items representing the same movement aligns with prior work ^43^ indicating that movement-specific variance can influence factor structures beyond modality distinctions in imagery questionnaires. This suggests that some movements are inherently more similar across visual and kinesthetic imagery, highlighting the importance of movement-specificity when interpreting questionnaires that rely on single movements (MIQ), daily movements (VMIQ), or sequential whole-body sports-related movements (e.g., ski-jumping).

The MIQ-RS had moderate to good test-retest reliability, consistent with prior laboratory-based studies ^13,56,57^ and other similar instruments ^12,58^, supporting its stability as a trait measure of movement imagery ability. Notably, the kinesthetic subscale showed higher reliability than the visual subscale (as opposed to the original validation study ^13^), suggesting that participants may provide more consistent ratings of kinesthetic sensations over time, whereas visual imagery may be more variable across sessions. This difference aligns with previous reports ^12^ indicating that kinesthetic imagery reflects more stable, body-centred representations, while visual imagery can be influenced by context or individual strategies.

Overall, our findings support the MIQ-RS as a robust tool for online assessment and reinforce the suitability of the MIQ-RS for longitudinal within-participants assessments and interventions, particularly when kinesthetic imagery is of primary interest.

### iFST

At the group level, execution times differed reliably between simple and complex sequences, and this effect was also present, though reduced, during imagery trials, indicating that the task captures graded differences in sequence difficulty consistent with motor planning processes ^15^. The iFST demonstrated strong internal validity in an online sample, extending previous laboratory-based studies of imagined movement sequences ^59,60^. These findings align with previous work showing that imagined and executed movements share temporal characteristics ^39,61^, supporting the construct validity of the iFST for assessing movement imagery maintenance. However, reliability analyses at the individual level only indicated moderate to good consistency across trials, suggesting that more trials may have been necessary (10 trials per condition were collected in this study). Additionally, the main effects and their interaction (illustrating the extent to which the constraint of sequence complexity was different in execution and imagery) could have shown greater reliability. This could indicate that the current paradigm might not be fully accurate to capture subtle differences in imagery ability. However, collecting more trials increases the likelihood of observing learning effects, and therefore both aspects should be considered in future iterations of this new task.

The iFST demonstrated moderate to good test-retest reliability for traditional mental chronometry measures, with both the absolute and relative difference scores showing consistent performance across sessions. These findings indicate that individuals’ temporal accuracy in imagery relative to execution was reasonably stable, supporting the utility of these measures for assessing movement imagery ability ^16,17^. In contrast, the Constraint score showed poor reliability (*ICC* = 0.33), suggesting that proportional indices capturing how well imagery preserves task-specific constraints may be more sensitive to intra-individual variability or trial-specific noise. This likely reflects the added complexity of the score, which involves differences in execution and imagery times across simple and complex sequences. Consequently, while mental chronometry remains a standard approach for capturing general imagery performance, the data suggest that the constraint score may be less suitable for individual-level assessments or longitudinal studies unless reliability can be improved through increased trial numbers or refined scoring procedures.

Overall, our findings support the iFST as a tool for assessing movement imagery in online samples and highlight the suitability of its traditional mental chronometry measures for longitudinal assessments and interventions, while cautioning that constraint-based indices may require refinement to achieve reliable individual-level measurement.

### HLJT

The HLJT demonstrated strong internal validity in an online setting, replicating well-established behavioural effects ^62–64^. Accuracy and reaction time differed reliably depending on rotation angle and hand view, with lateral rotations processed less accurately and more slowly than medial rotations, particularly in the palmar view, reflecting the influence of biomechanical constraints. The ‘grand’ biomechanical constraints effect further confirmed that imagined movements are sensitive to anatomical limitations at the group level, with larger differences observed in the palmar view compared to the dorsal view ^65^. Reliability analyses indicated moderate consistency across trials, suggesting that, while the HLJT provides robust group-level effects, individual-level measures may be more variable. This highlights the necessary use of individual-level analyses considering trial-level variability in this task.

The HLJT demonstrated good test-retest reliability, with both accuracy, reaction time and biomechanical constraints showing stable data across sessions. These findings, which are in line with previous work ^25^, indicate that participants’ performance is consistent over time, supporting the suitability of the HLJT for longitudinal assessments. Combined with the good internal validity of fundamental behavioural effects, the HLJT appears to provide reliable measures of the cognitive representation of hand movements, making it a valuable tool for both experimental and applied settings.

Overall, these results support the HLJT as a valid and reliable tool for assessing movement imagery in online settings, reinforcing its suitability for large-scale or longitudinal studies investigating the cognitive representation of hand movements.

### Relationships between tests

Overall, the results suggested that performance across the different processes of movement imagery is largely independent, consistent with a substantial body of previous evidence ^6,66,67^. Bayesian correlations indicated that most measures did not correlate, although a few weak-to-negligible associations were observed. These findings imply that performance in one process of movement imagery (e.g., generation, maintenance, or manipulation) does not reliably predict performance in another, emphasizing that individual differences are process- and task-specific. This underscores the necessity of assessing multiple processes when evaluating imagery ability and tailoring the choice of measures to the context—whether for research purposes or applied interventions—since relying on a single metric may fail to capture the full spectrum of an individual’s imagery skills ^9^.

The reasons why self-report measures (MIQ-RS) and behavioural measures (iFST, HLJT) do not correlate are still a matter of debate. First, imagery questionnaires ask individuals to objectify the quality of an experience, without having experienced the possible range of qualities before^68^. When completing imagery questionnaires, individuals are subject to avoidance of negative self-image ^69,70^ and a number of other biases (including response biases) ^71^. While it has been suggested that self-reports are better measurement instruments than behavioural tasks ^72^, the importance of behavioural measures in neuroscience is clear ^73^. These two types of measures certainly tap into different aspects of movement imagery and may or may not agree when it comes to assessing imagery ability.

The lack of correlation between behavioural tasks (iFST and HLJT) may be because each task truly taps into distinct imagery-related processes, despite being broadly categorised under movement imagery. For instance, the iFST requires participants to maintain a motor sequence over time, engaging processes related to working memory, temporal stability of imagery, and possibly motor planning ^60,74^. In contrast, the HLJT assesses the ability to mentally rotate hand images, relying more heavily on visuospatial transformation and perspective-taking ^26^. These tasks, although both categorised under movement imagery, likely recruit different neural and cognitive systems. Additionally, task-specific factors such as instructions, response modalities, and contextual demands may introduce variability ^34^. Together, these considerations suggest that behavioural tasks, while valuable, may not be interchangeable proxies for a unified imagery ability, but rather complementary tools that assess different processes of movement imagery.

Collectively, our results highlight the value of a multidimensional, process-specific approach for both experimental and applied assessments of movement imagery. More broadly, these results also highlight the need for studies with larger samples able to detect small effects. Future studies should investigate whether different behavioural tasks aiming to assess a specific imagery process would show meaningful correlations, as this would provide evidence that they are indeed measuring the same underlying skill.

### Strengths and limitations

This study is among the first to propose three tests targeting the generation, manipulation, and maintenance of movement imagery ability in an online setting, using robust procedures and validated measures. A limitation is that the iFST may not have included enough trials per condition to yield reliable estimates, though this can be easily addressed in future iterations. A potential learning effect should also be considered given that the test-retest period was only one week, though results from ICCs and measurement error metrics showed this was negligible.

Another limitation is that each domain was assessed with a single task, making it difficult to determine whether the lack of correlations reflects true independence between processes or simply task-specific differences. However, instruments thought to measure the same construct sometimes do not converge well if they differ in measurement method (questionnaire vs. behavioural task) ^75^ or dependent variables (ratings vs. reaction time vs. error rates).

Nonetheless, the study successfully achieved its goal of identifying and validating targeted measures in a fully online setting, providing a basis for future research to expand and refine this approach. Finally, the time to complete each of these tests varied between 15-20 minutes. This limits the practical implementation of all tests into other experiments. However, we recommend applied researchers to consider which specific process of imagery is more relevant to their design, in case they need to optimise testing, or consider developing shorter versions of these assessments in future studies.

## CONCLUSIONS

This study provides evidence supporting the validity and reliability of an online resource to assess movement imagery ability, encompassing the MIQ-RS, iFST, and HLJT for generation, maintenance and manipulation. The MIQ-RS demonstrated robust structural validity, adequate internal consistency, and moderate to good test-retest reliability, particularly for the kinesthetic subscale. The iFST showed strong internal validity, capturing graded differences in sequence difficulty, with moderate test-retest reliability for traditional mental chronometry measures, though constraint-based indices were less stable. The HLJT effectively reproduced well-established stimulus rotation, hand view and biomechanical constraints effects and exhibited good test-retest reliability for both accuracy and reaction time. Importantly, correlations between tasks were generally weak or negligible, indicating that performance in one imagery process does not reliably predict performance in another. This emphasizes the need for multidimensional assessment when evaluating movement imagery ability and for tailoring task selection to the specific research or applied context. Collectively, these findings highlight the utility of combining self-report and behavioural measures to capture the full spectrum of imagery skills. The results underscore the importance of large, well-powered studies to further clarify subtle interrelations across imagery domains and refine multidimensional assessment frameworks for both experimental and applied applications.

## ACKNOWLEDGEMENTS

We would like to thank Andrés de los Santos for his help in developing the figures. We would also like to thank Álvaro Gonzalo, Guillermo García-Hidalgo, Diego Caso, Marta Figueroa, Sara Lobato, Antonio Pavón, Almudena Cerezo, Concepción Verdú and Sonia Moreno for their help in the piloting phase.

## ETHICS APPROVAL STATEMENT

Ethical approval was obtained by the Ethics Commission of the Institute for Research in the Psychological Sciences at UCLouvain, Belgium (ID: Projet2024-85). The study was performed in accordance with the ethical standards as laid down in the 1964 Declaration of Helsinki and its later amendments.

## CONSENT TO PARTICIPATE STATEMENT

All individuals gave explicit electronic informed consent before participating.

## CONSENT FOR PUBLICATION STATEMENT

All participants provided electronic informed consent for the anonymised data collected during the study to be used in scientific publications. All participants also agreed that a fully anonymised dataset was to be deposited in an Open Data Repository at the end of the project.

## CODE AVAILABILITY STATEMENT

The code and materials for all experiments are available at https://osf.io/7cfz4/.

## OPEN PRACTICES STATEMENT

The data and code used for data analysis are available at https://osf.io/7cfz4/.

## AUTHOR CONTRIBUTIONS

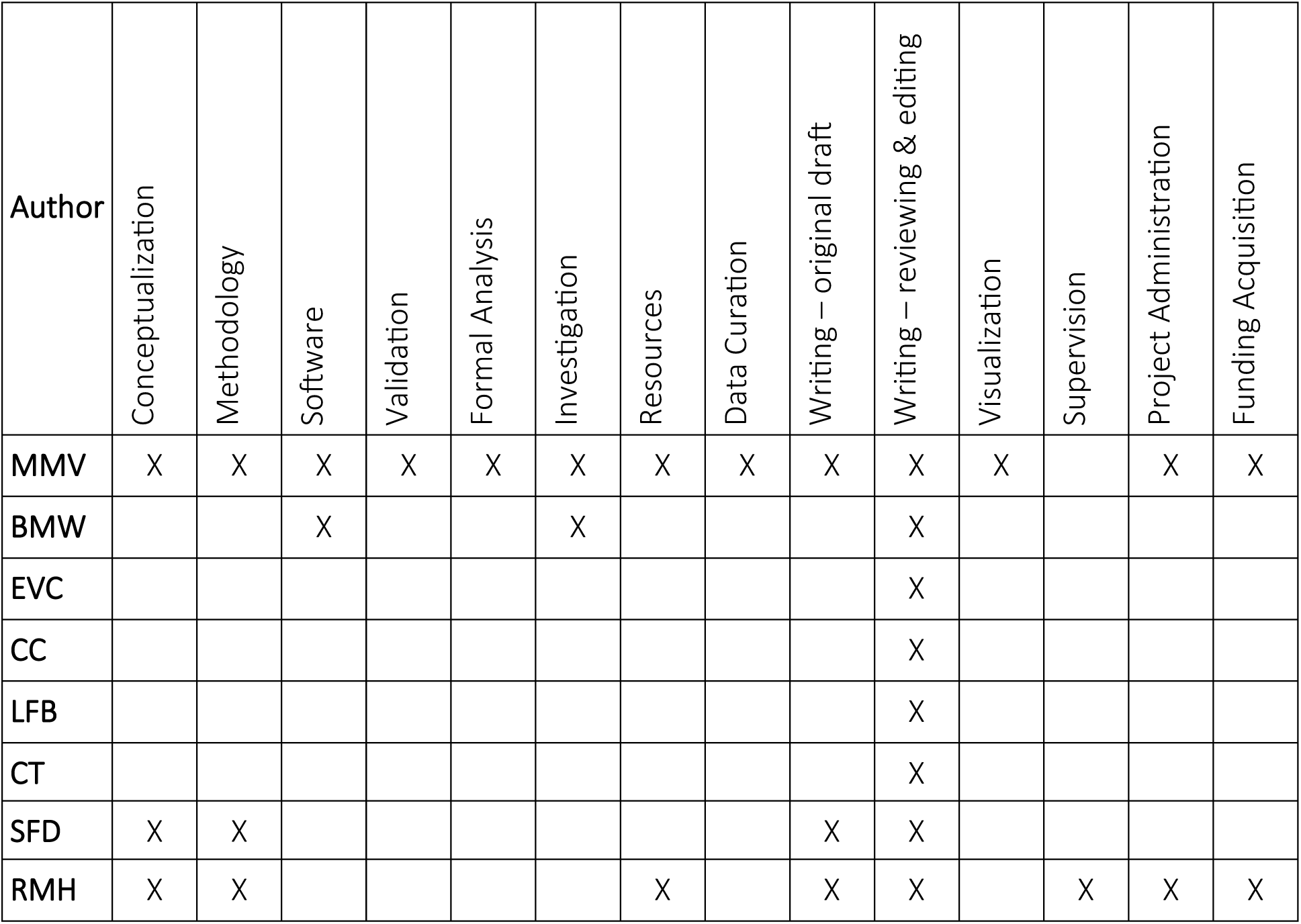

