## Supplementary Materials for "A comprehensive, open-source battery of movement imagery ability tests: Development and psychometric properties"

### METHODS

#### Assessment of levels of tiredness/fatigue and attention

The level of tiredness/fatigue and attention was self-reported before the Test and Retest session on 11-point scales. For tiredness/fatigue, the scale went from 0 = not tired/fatigued at all to 10 = very tired/fatigued. For attention, the scale went from 0 = extremely difficult to focus to 10 = extremely easy to focus. Participants reporting an absolute difference between Test and Retest > 2 points were excluded in the sensitivity analysis of test-retest reliability.

#### Self-assessment questions

All these questions were reported descriptively without any pre-planned statistical analysis.

<u>MIQ-RS:</u> Participants were asked which modality (visual or kinesthetic) was their preferred (0 = Strong preference visual, to 10 = Strong preference kinesthetic). They were also asked, when they imagined in a visual modality, which perspective were they using (0 = Totally first-person, to 10 = Totally third-person). Finally, they were asked about possible involuntary use of the visual/kinesthetic modalities when instructed to use kinesthetic/visual modalities (0 = No visual/kinesthetic use, to 10 = High visual/kinesthetic use).

<u>iFST:</u> Participants were asked which was their use of visual and kinesthetic modalities during imagery in each block of trials, that is for each sequence type separately (0 = No kinesthetic/visual imagery use, to 10 = High kinesthetic/visual imagery use). They were also asked which visual perspective (first-person or third-person) they employed, if they used a visual modality (0 = No use, to 10 = High use). Additionally, they were asked whether they imagined themselves making errors during imagery (0 = No errors, I was completely accurate, to 10 = A lot of errors, I was completely inaccurate). Finally, they were asked about their overall vividness (0 = No image at all, you only "know" that you were thinking of the movement, to 10 = Perfectly clear and vivid as normal vision and/or feel of movement).

<u>HLJT:</u> Participants were first asked through an open question to briefly describe which strategies they used throughout the task. Then, they were asked whether they "thought about" their own hands during the task (yes/no), and if so, to briefly explain what they did. Afterwards, they were explicitly asked whether they were imagining moving their hands to make the laterality judgements (yes/no) and if so, which hand were they imagining (left, both (sometimes left, sometimes right), or right). After that, they were asked if they used visual/kinesthetic imagery (0 = No kinesthetic/visual imagery use, to 10 = High kinesthetic/visual imagery use) and which visual perspective, first-person or third-person (0 = No use, to 10 = High use).

### Statistical Analysis details

Generative Hierarchical Bayesian Models: The models were created mimicking the frequentist GLMMs of the internal validity analysis. However, we had to include some technical modifications to these Bayesian models, which are explained below.

- The formula of the models included not only the fixed effects parameters (μ 'submodel') but also parameters for the variability of the effects (σ 'submodel'). We allowed correlations between the random effects (intercepts and slopes).
- The models were run with 5 chains, 10,000 iterations per chain and 2,000 warmup iterations per chain, for a total post-warmup of 40,000 iterations. Effective sample sizes (ESS) and $R^2$ values were checked alongside posterior predictive checks to assess goodness-of-fit.
- The models used "regularizing" priors in the sense that the priors over the beta parameters were non-informative but restricted implausible coefficients.
    - For the iFST, we used a prior of a normal distribution with mean = 0 and SD = 3 for the fixed effects in the mean (μ) 'submodel', reflecting a wide belief that the effects of trial type, sequence complexity, or their interaction could plausibly reach up to ±3 seconds, but are otherwise weakly informative. For the intercept, we used a normal distribution with mean = log(3) and SD = 1, corresponding to an expected baseline time to complete a sequence of approximately 3 seconds with moderate uncertainty (based on the frequentist GLMM's results). For the standard deviations of the random effects (intercepts and slopes for each participant), we used an exponential prior with rate 1, and for the correlations between random effects we applied an LKJ prior with parameter 2. Finally, for the fixed effects on the residual standard deviation (σ 'submodel'), we used a normal prior with mean = 0 and SD = 1, which is somewhat conservative. All priors on the fixed effects are weakly informative, providing regularization without strongly constraining the estimates.
    - For the HLJT, we used a prior of a normal distribution with mean = 0 and SD = 0.5 for the fixed effects in the mean (μ) 'submodel'. This reflects the expectation that predictors are unlikely to shift the log-reaction times by more than about ±1.5 units on the log scale, which corresponds to moderate changes on the millisecond scale, while remaining weakly informative. For the intercept, we used a prior of a normal distribution with mean = log(1500) and SD = 0.5, corresponding to the expectation that the typical reaction time should be around 1500ms, with moderate uncertainty. For the variance (σ) 'submodel', we used a prior of a normal distribution with mean = 0 and SD = 0.5 for the effects of predictors on log(σ), and a normal distribution with mean = 0 and SD = 1 for the intercept of log(σ). Together, these priors are regularizing, constraining the model to plausible values without being strongly informative.
- Signal-to-noise ratios (SNRs) were computed by extracting random effects coefficients directly from the model object. For each participant, the signal was the average of the effect of a parameter and the noise the SD of that effect. To calculate the group-level SNR (reported in Table 3 of the main manuscript), we divided the SD of the signal by the average of the noise ($SNR = \sigma_{signal} / \mu_{noise}$). Specifically, we computed the standard

deviation of the posterior means of each participant's effect average estimates (signal) and divided it by the mean of the posterior standard deviation of trial-level data (noise). This provides a robust group-level index of reliability that incorporates both between- and within-participant variability, aligning with recent approaches to quantify consistency in individual-level estimates in hierarchical models.

Learning effect in the iFST: We ran an analysis to rule out the possibility that the relatively limited reliability in the iFST was due to participants improving their performance (i.e. reducing the difference between execution and imagery) by just repeating the execution/imagination of the sequence multiple times (illustrating a learning effect). As each participant completed a maximum of 10 trials, we computed the differences between imagery times (each trial) with the average execution time (across all trials), for each sequence. We then created two trial groups: the first 5 trials and the last 5 trials and ran linear mixed models considering the sequence type and the trial group as predictors. We were interested in the interaction between these two factors, whereby the last few trials would show smaller differences between execution and imagery compared to the last trials, for a specific sequence type.

Sensitivity analyses test-retest reliability and measurement error: All analyses were repeated excluding participants whose difference between Test and Retest in tiredness/attention or level of attention exceeded 2 points on the 11-point scale.

#### Sample size calculations

All calculations below were conducted with 5% Type I error rate and 80% power. Effect sizes were considered small-to-moderate. Adjustments for multiple comparisons were done via Bonferroni correction. We decided to collect data from $N$ = 180 participants for the Test session, as it would be powered for all our pre-planned analyses.

For structural validity of the MIQ-RS, a power calculation for Confirmatory Factor Analysis was conducted. With an expected factor correlation $r$ = 0.3 and an average factor loading of $r$ = 0.5, both of which are conservative based on previous reports from the MIQ-RS questionnaire [1,2], indicated $N$ = 176 participants were needed (calculated with 'semPower' package v2.1.1 [3]).

For internal validity of behavioural tasks pre-planned post-hoc paired t-tests with two tails, assuming a weak-to-moderate effect size of Cohen's d = 0.3 were used (calculated with 'pwr' package v1.3-0 in R [4]). $N$ = 108 participants would be needed for the iFST because we would perform 2 tests, simple vs complex sequence times for 2 trial types, therefore the Bonferroni-corrected p-value would be $p$ = 0.05/2 = 0.025. For the HLJT, $N$ = 145 participants would be needed as we would perform 8 tests in the strictest post-hoc scenario, in the case of 8 rotation angles tested sequentially, the corrected p-value would be $p$ = 0.05/8 = 0.006.

For criterion validity, a weak correlation coefficient of $r$ = 0.3 in a two-tailed contrast and adjusting for multiple comparisons in the correlation matrix with 15 tests of interest (2 MIQ-RS measures, 3 iFST measures, 2 HLJT measures, each measure correlated with measures of the other tests, but not with measures within their own test) was used. $N$ = 152 participants would be needed for a Bonferroni-corrected p-value of $p$ = 0.05/15 = 0.003 (calculated with 'pwr' package).

RESULTS

**Supplementary Figure 1.** Flowchart of participants throughout data collection steps. Criteria by task: 1) Movement Imagery Questionnaire-Revised Second edition (MIQ-RS): 1 attention check half-way through the questionnaire. 2) Imagined Finger Sequence Task (iFST): >75% accuracy in execution & imagery AND average execution & imagery time >1 second. 3) Hand Laterality Judgement Task (HLJT): >60% accuracy overall & >75% trials available for analysis (reaction time > 300ms and < 3,000ms).

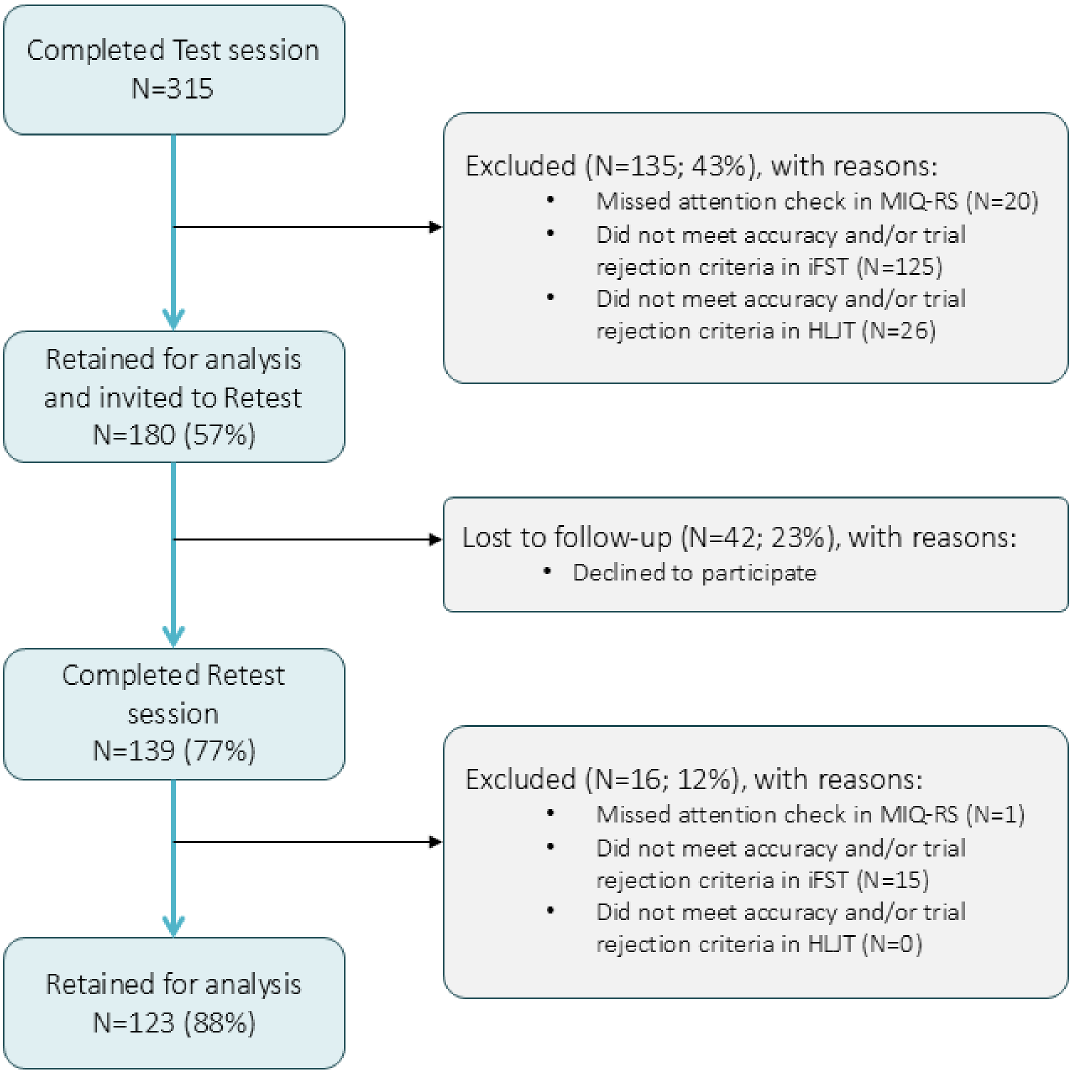

**Supplementary Table 1.** Item-total correlations and correlations if item removed of the Movement Imagery Questionnaire-Revised Second edition (MIQ-RS).

| Item | Subscale | Item-total correlation | Correlation if removed |
|---|---|---|---|
| 1 | Kinesthetic | 0.646 | 0.609 |
| 2 | Visual | 0.465 | 0.420 |
| 3 | Kinesthetic | 0.615 | 0.574 |
| 4 | Visual | 0.501 | 0.441 |
| 5 | Visual | 0.445 | 0.391 |
| 6 | Kinesthetic | 0.601 | 0.544 |
| 7 | Kinesthetic | 0.635 | 0.585 |
| 8 | Visual | 0.585 | 0.526 |
| 9 | Kinesthetic | 0.637 | 0.593 |
| 10 | Visual | 0.558 | 0.495 |
| 11 | Kinesthetic | 0.591 | 0.549 |
| 12 | Kinesthetic | 0.600 | 0.553 |
| 13 | Visual | 0.609 | 0.547 |
| 14 | Visual | 0.594 | 0.531 |

**Supplementary Figure 2.** Distributions of subscale sum-scores in the Movement Imagery Questionnaire-Revised Second edition (MIQ-RS).

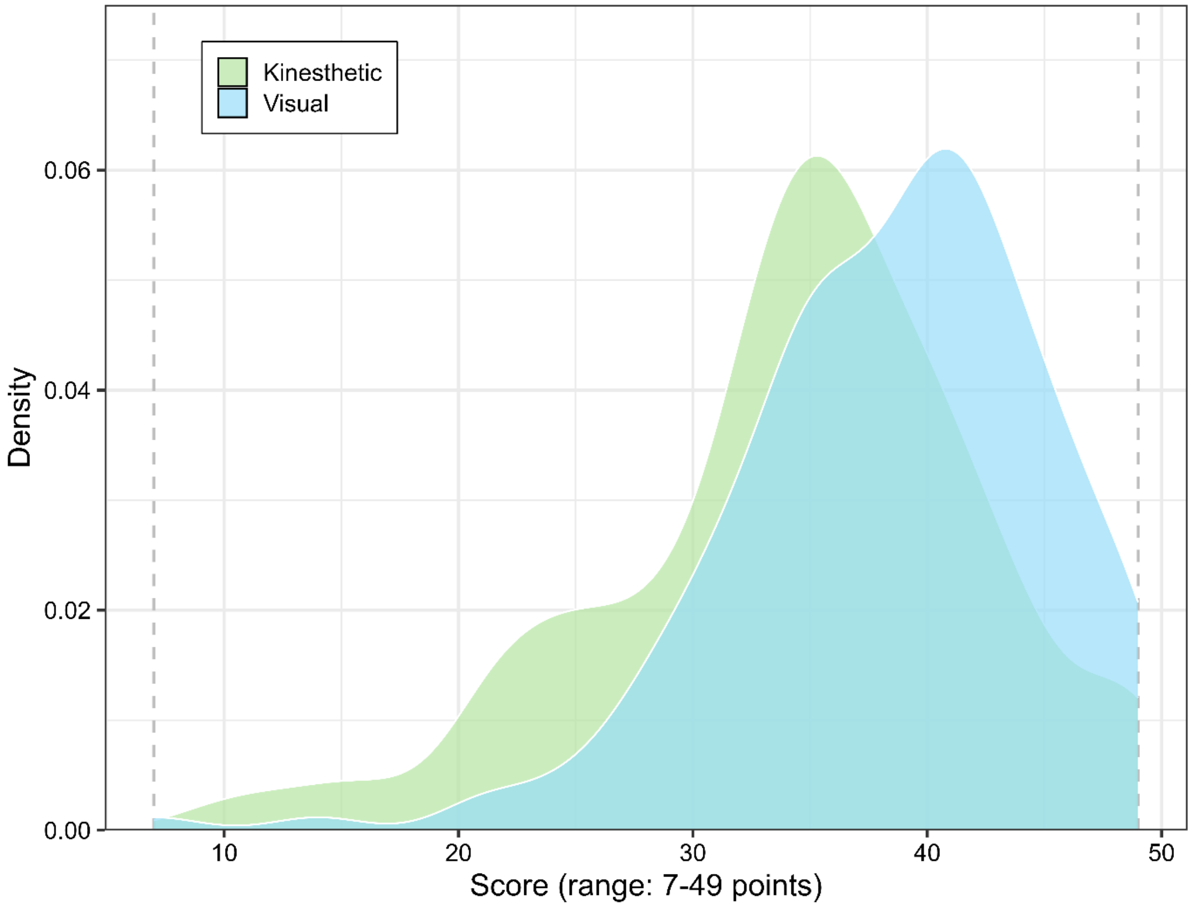

**Supplementary Figure 3.** Reliability in the Imagined Finger Sequence Task (iFST). Predictions from the generative hierarchical Bayesian model for time to complete a sequence (seconds), according to each condition in the task (sequence complexity: visual vs complex; trial type: execution vs imagery; and their interaction), and the intercept. The plot shows the posterior mean and 95% credible interval for each individual (N=180) for each condition, illustrating within-participant variability (width of credible interval) and between-participant variability (dispersion of means across participants), which together illustrate the noise and signal in the task, respectively. Data are back-transformed from the log scale, therefore the intercept is on the original metric (seconds) but the fixed effects are on a multiplicative metric (meaning that, for example, a value of 1.2 would indicate a difference of 20% between the levels of that factor, and a value of 1 would be a zero difference).

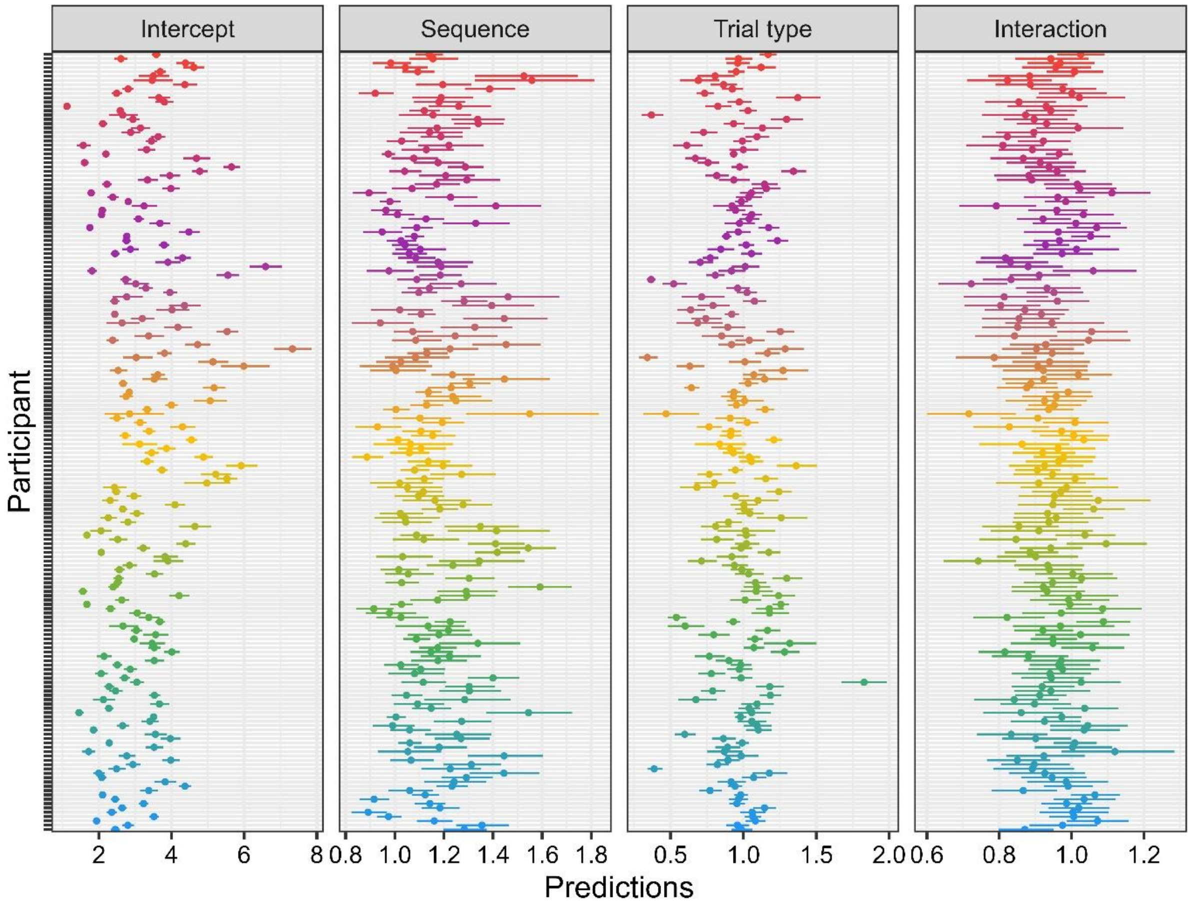

### Sensitivity Analysis iFST validity and reliability

After excluding participants missing at least 1 out of 2 instruction checks in the iFST ($N$ = 52), the results did not meaningfully change. The GLMM included $N$ = 128 participants and 4779 trials. For execution, the model estimated an average of 3.00 seconds [2.76, 3.24] to complete the simple sequence and 3.92 seconds [3.55, 4.28] to complete the complex sequence, with evidence for a difference between the two (post-hoc mean difference = -0.91 seconds [-1.15, -0.68], $t_{(4764)}$ = -7.59, $p$ = 7.68x$10^{-14}$). For imagery, the average time to complete the simple sequence was 3.00 seconds [2.7, 3.29] and for the complex sequence 3.36 seconds [3.04, 3.69], with evidence for a difference but with a smaller effect size (post-hoc mean difference = -0.36 seconds [-0.57, -0.16], $t_{(4764)}$ = -5.21, $p$ = 3.88x$10^{-7}$). These results are largely equivalent to the ones obtained through the model including $N$ = 180 participants.

Regarding reliability, the results for the SNRs were equivalent to the main analysis including the full sample: Intercept = 8.81, Trial type = 4.92, Sequence type = 2.82, Interaction = 1.57.

### Learning effects in the iFST

The analysis revealed that for the simple sequence, both the raw difference and the relative difference where statistically lower for the last 5 trials compared to the first 5 trials (raw mean difference (first vs last) = -0.08 seconds [-0.14, -0.1], $t_{(3320)}$ = -2.3, $p$ = 0.021; relative mean difference (first vs last) = -3.15% [-5.33, -1.07], $t_{(3320)}$ = -2.97, $p$ = 0.003). However, this did not occur for the complex sequence (raw mean difference (first vs last) = -0.03 seconds [-0.12, 0.06], $t_{(3320)}$ = -0.66, $p$ = 0.507; relative mean difference (first vs last) = -1.68% [-3.93, 0.56], $t_{(3320)}$ = -1.47, $p$ = 0.141). These results illustrate practically minimal (less than 0.1 seconds representing around 3% of error), though statistically significant, learning effects only in the simple sequence in the iFST.

**Supplementary Table 2.** Accuracy (%) contrasts with pairs of consecutive rotation angles in the Hand Laterality Judgement Task, for each hand view.

| View | Contrast | Difference (%) | 95% CI | Z statistic | p-value |
|---|---|---|---|---|---|
| Dorsal | 0° vs 45° | 0.04 | [-1.63, 1.72] | 0.05 | 0.961 |
| | 45° vs 90° | -2.06 | [-3.88, -0.23] | -2.21 | 0.027 |
| | 90° vs 135° | -7.48 | [-9.71, -5.25] | -6.58 | $4.76\times10^{-11}$ |
| | 135° vs 180° | -11.21 | [-14.07, -8.36] | -7.70 | $1.37\times10^{-14}$ |
| | 180° vs 225° | 13.59 | [10.79, 16.39] | 9.52 | $1.78\times10^{-21}$ |
| | 225° vs 270° | 6.02 | [3.92, 8.12] | 5.61 | $1.97\times10^{-8}$ |
| | 270° vs 315° | 2.64 | [0.82, 4.47] | 2.84 | 0.005 |
| Palmar | 0° vs 45° | -2.15 | [-3.86, -0.43] | -2.46 | 0.014 |
| | 45° vs 90° | -3.62 | [-5.73, -1.51] | -3.36 | $7.68\times10^{-4}$ |
| | 90° vs 135° | -7.11 | [-9.59, -4.64] | -5.63 | $1.82\times10^{-8}$ |
| | 135° vs 180° | 2.45 | [-0.4, 5.3] | 1.68 | 0.093 |
| | 180° vs 225° | 9.20 | [6.96, 11.45] | 8.04 | $8.92\times10^{-16}$ |
| | 225° vs 270° | 1.13 | [-0.47, 2.73] | 1.38 | 0.168 |
| | 270° vs 315° | 1.32 | [-0.11, 2.74] | 1.81 | 0.070 |

Note: Differences are taken subtracting highest minus lowest angle, therefore for contrasts <180° the differences are negative because the highest angle shows lower accuracy.

**Supplementary Table 3.** Reaction time (ms) contrasts with pairs of consecutive rotation angles in the Hand Laterality Judgement Task, for each hand view.

| View | Contrast | Difference (ms) | 95% CI | T statistic | p-value |
|---|---|---|---|---|---|
| Dorsal | 0° vs 45° | 55.76 | [26.3, 85.23] | 3.71 | $2.08\times10^{-4}$ |
| | 45° vs 90° | 116.86 | [88.86, 144.87] | 8.18 | $2.92\times10^{-16}$ |
| | 90° vs 135° | 162.75 | [130.48, 195.01] | 9.89 | $5\times10^{-23}$ |
| | 135° vs 180° | 213.59 | [161.17, 266.01] | 7.99 | $1.42\times10^{-15}$ |
| | 180° vs 225° | -247.98 | [-299.93, -196.02] | -9.36 | $8.7\times10^{-21}$ |
| | 225° vs 270° | -156.73 | [-187.48, -125.98] | -9.99 | $1.79\times10^{-23}$ |
| | 270° vs 315° | -134.13 | [-165.16, -103.09] | -8.47 | $2.52\times10^{-17}$ |
| Palmar | 0° vs 45° | 44.91 | [21.27, 68.55] | 3.72 | $1.96\times10^{-4}$ |
| | 45° vs 90° | 115.03 | [81.94, 148.12] | 6.81 | $9.65\times10^{-12}$ |
| | 90° vs 135° | 146.47 | [106.88, 186.05] | 7.25 | $4.18\times10^{-13}$ |
| | 135° vs 180° | -26.57 | [-73.94, 20.79] | -1.10 | 0.272 |
| | 180° vs 225° | -236.68 | [-274.23, -199.13] | -12.35 | $5.24\times10^{-35}$ |
| | 225° vs 270° | -91.47 | [-115.73, -67.2] | -7.39 | $1.5\times10^{-13}$ |
| | 270° vs 315° | -27.57 | [-50.78, -4.36] | -2.33 | 0.020 |

Note: Differences are taken subtracting highest minus lowest angle, therefore for contrasts >180° the differences are negative because the highest angle shows shorter reaction time.

Supplementary Figure 4. Reliability in Hand Laterality Judgement Task (HLJT). Predictions from the generative hierarchical Bayesian model for Reaction Time (ms), according to each condition in the task (direction: medial or lateral; hand view: palmar and dorsal; and their interaction), and the intercept. The plot shows the posterior distribution for each individual (N=180) for each condition, illustrating participant-level variability and condition-level variability, which together illustrate the signal-to-noise ratio. Data are back-transformed from the log scale, therefore the intercept is on the original metric (milliseconds) but the fixed effects are on a multiplicative metric (meaning that, for example, a value of 1.2 would indicate a difference of 20% between the levels of that factor, and a value of 1 would be a zero difference).

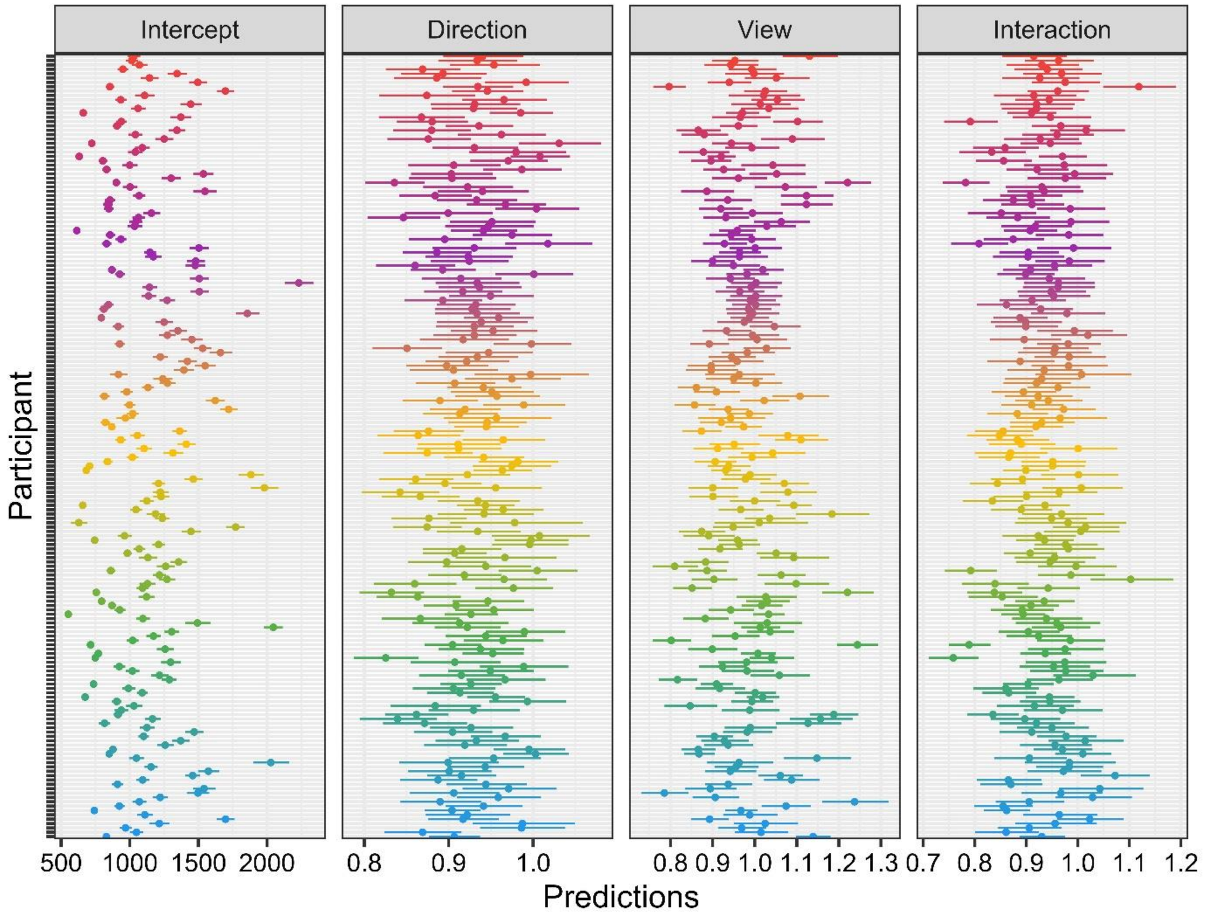

**Supplementary Table 4.** Criterion validity analysis according to Spearman's rank correlation coefficients in a Bayesian framework.

| Variable 1 | Variable 2 | Spearman rho | 95% CrI | Bayes Factor |
|---|---|---|---|---|
| MIQ-RS Visual | MIQ-RS Kinesthetic | 0.37 | [0.25, 0.5] | **$BF_{10}$ = 1.09×10$^{5}$** |
| MIQ-RS Visual | iFST Difference (raw) | -0.15 | [-0.29, -0.01] | **$BF_{10}$ = 1.07** |
| MIQ-RS Visual | iFST Difference (relative) | -0.13 | [-0.27, 0.02] | $BF_{01}$ = 1.83 |
| MIQ-RS Visual | iFST Constraint score | -0.08 | [-0.22, 0.07] | $BF_{01}$ = 4.02 |
| MIQ-RS Visual | HLJT Accuracy | 0.12 | [-0.01, 0.27] | $BF_{01}$ = 1.93 |
| MIQ-RS Visual | HLJT Reaction Time | 0.00 | [-0.15, 0.14] | $BF_{01}$ = 7.76 |
| MIQ-RS Kinesthetic | iFST Difference (raw) | -0.02 | [-0.17, 0.12] | $BF_{01}$ = 7.27 |
| MIQ-RS Kinesthetic | iFST Difference (relative) | -0.04 | [-0.17, 0.12] | $BF_{01}$ = 6.98 |
| MIQ-RS Kinesthetic | iFST Constraint score | -0.05 | [-0.2, 0.09] | $BF_{01}$ = 6 |
| MIQ-RS Kinesthetic | HLJT Accuracy | 0.17 | [0.02, 0.3] | **$BF_{10}$ = 1.72** |
| MIQ-RS Kinesthetic | HLJT Reaction Time | 0.03 | [-0.11, 0.18] | $BF_{01}$ = 7.12 |
| iFST Difference (raw) | iFST Difference (relative) | 0.91 | [0.89, 0.94] | **$BF_{10}$ = 4.03×10$^{69}$** |
| iFST Difference (raw) | iFST Constraint score | 0.66 | [0.57, 0.74] | **$BF_{10}$ = 3.11×10$^{21}$** |
| iFST Difference (raw) | HLJT Accuracy | -0.11 | [-0.25, 0.03] | $BF_{01}$ = 2.47 |
| iFST Difference (raw) | HLJT Reaction Time | 0.27 | [0.14, 0.4] | **$BF_{10}$ = 141.32** |
| iFST Difference (relative) | iFST Constraint score | 0.57 | [0.46, 0.66] | **$BF_{10}$ = 1.61×10$^{14}$** |
| iFST Difference (relative) | HLJT Accuracy | -0.14 | [-0.28, 0.01] | $BF_{01}$ = 1.24 |
| iFST Difference (relative) | HLJT Reaction Time | 0.18 | [0.04, 0.31] | **$BF_{10}$ = 2.34** |
| iFST Constraint score | HLJT Accuracy | -0.05 | [-0.19, 0.09] | $BF_{01}$ = 6.11 |
| iFST Constraint score | HLJT Reaction Time | 0.13 | [-0.01, 0.27] | $BF_{01}$ = 1.46 |
| HLJT Accuracy | HLJT Reaction Time | 0.04 | [-0.1, 0.18] | $BF_{01}$ = 6.9 |

Note: Bayes Factors are presented in favour of the alternative hypothesis ($BF_{10}$, in bold) or the null hypothesis ($BF_{01}$). Abbreviations: 95% CrI: 95% Credible Interval; HLJT: Hand Laterality Judgement Task; iFST: Imagined Finger Sequence Task; MIQ-RS: Movement Imagery Questionnaire-Revised Second edition.

**Supplementary Figure 5.** Test-retest reliability of the Movement Imagery Questionnaire-Revised Second edition (MIQ-RS). **Panels a)** and **b)** show change from Test to Retest sessions (6-8 days apart), for visual and kinesthetic subscale sum-scores, respectively. **Panels c)** and **d)** show differences between Test and Retest plotted against the means as Bland-Altman plots. In these plots, dashed lines indicate the Limits of Agreement and solid line indicates mean difference (i.e., bias), with its 95% confidence interval marked in grey.

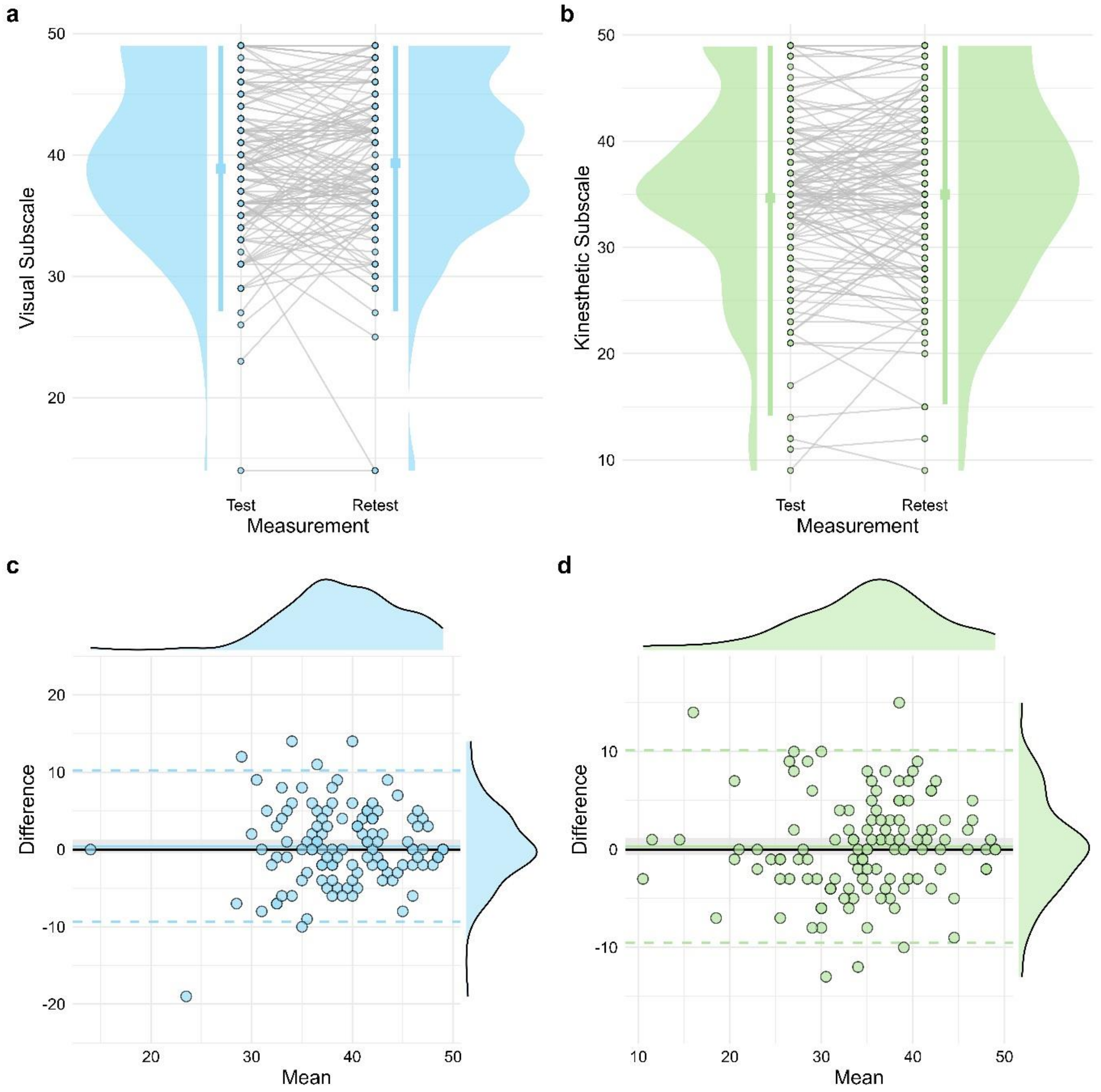

**Supplementary Figure 6.** Test-retest reliability of the Imagined Finger Sequence Task (iFST). **Panels a-c** show change from Test to Retest sessions (6-8 days apart), for Raw Difference, Relative Difference and Constraint Score, respectively. **Panels d-f** show differences between Test and Retest plotted against the means as Bland-Altman plots. In these plots, dashed lines indicate the Limits of Agreement and solid line indicates mean difference (i.e., bias), with its 95% confidence interval marked in grey.

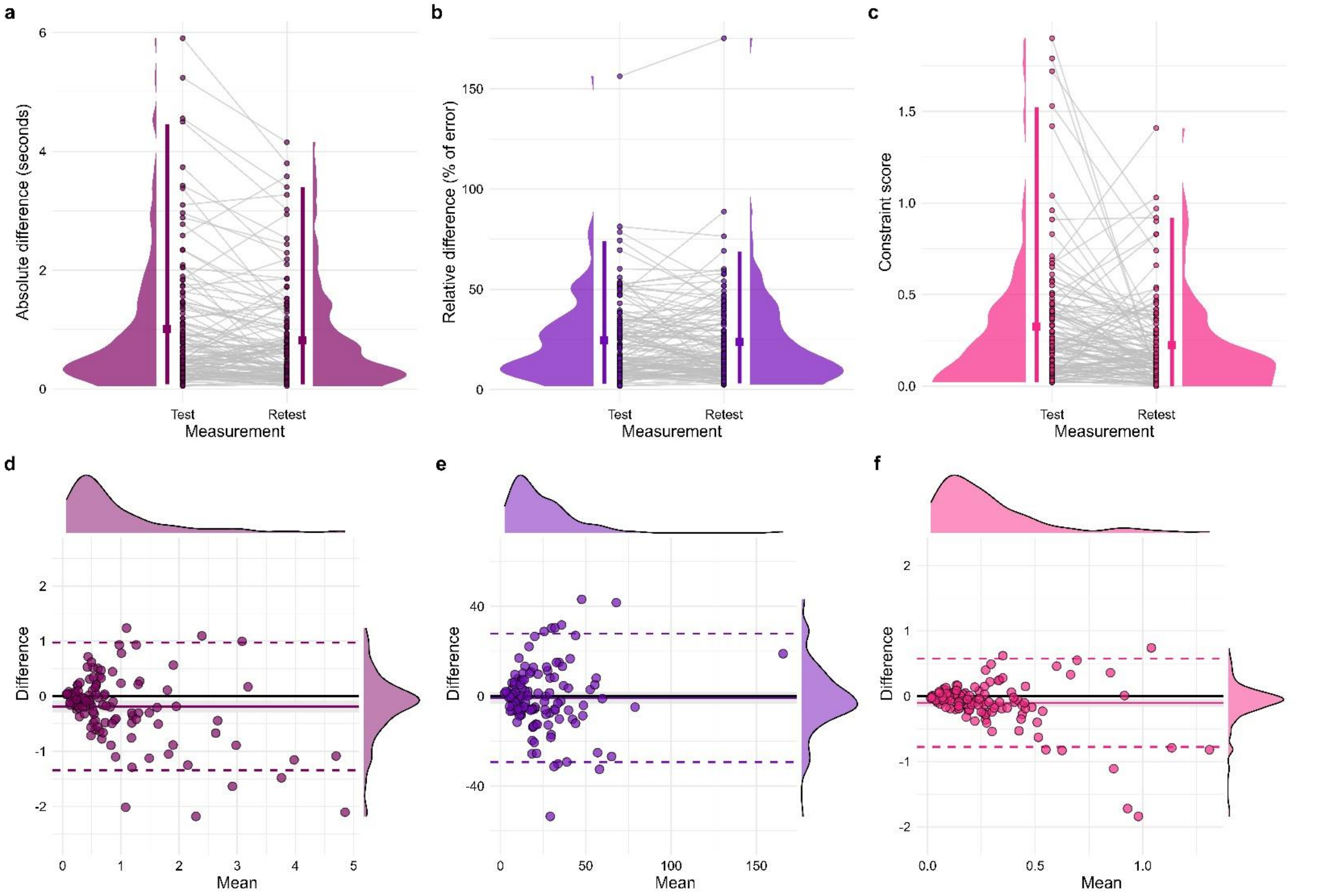

**Supplementary Figure 7.** Test-retest reliability of the Hand Laterality Judgement Task (HLJT). **Panels a-c** show change from Test to Retest sessions (6-8 days apart), for Accuracy, Reaction Time and Biomechanical Constraints, respectively. **Panels d-f** show differences between Test and Retest plotted against the means as Bland-Altman plots. In these plots, dashed lines indicate the Limits of Agreement and solid line indicates mean difference (i.e., bias), with its 95% confidence interval marked in grey.

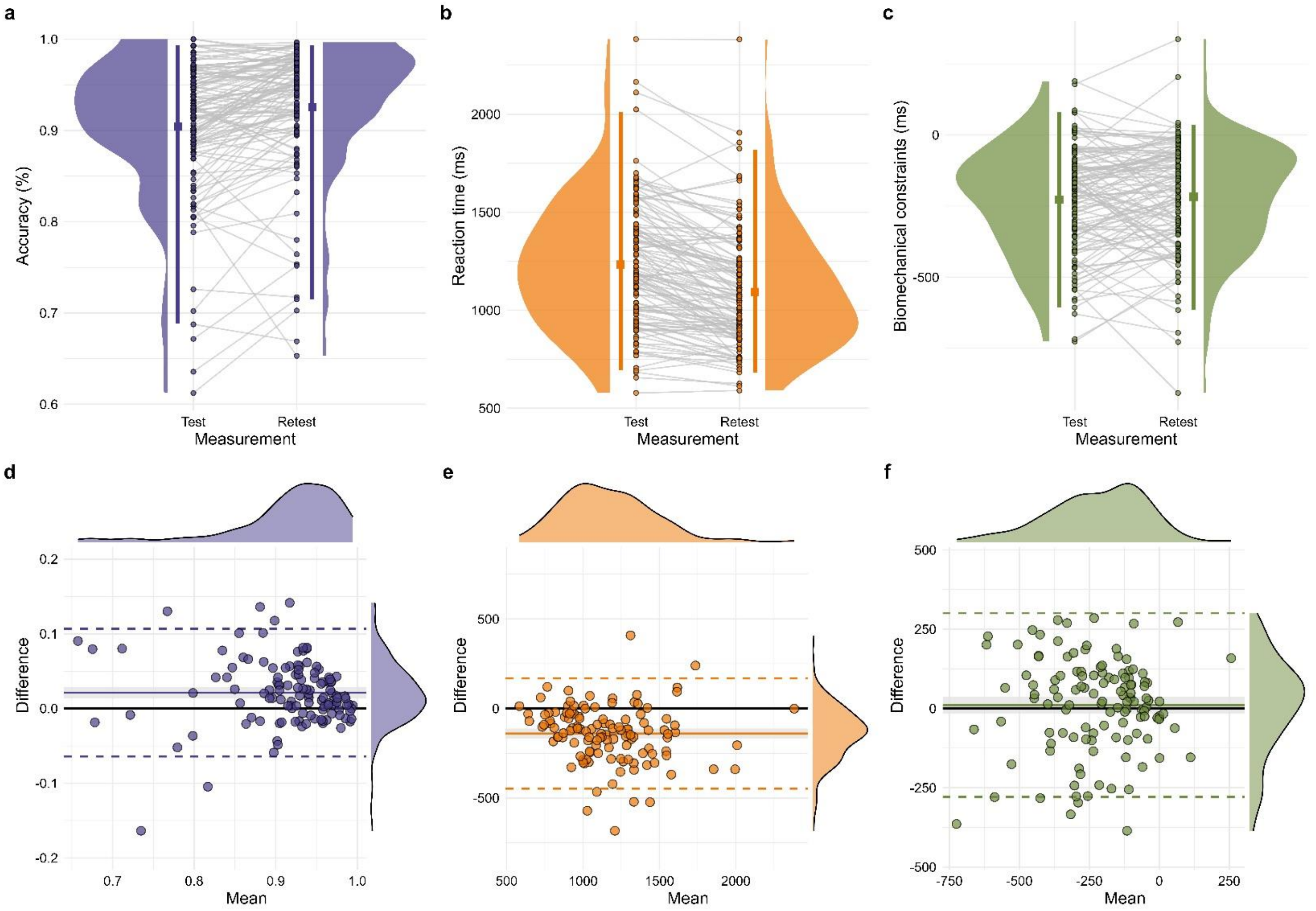

**Supplementary Table 5.** Sensitivity analysis for test-retest reliability, removing participants who had a difference >2 points (out of 10) in level of tiredness/fatigue and/or attention from Test to Retest (*N* = 22). Total participants analysed were *N* = 101.

| Test | Variable | Mean Test | SD Test | Mean Retest | SD Retest | ICC (2,1) | 95%CI | SEM | $MDC_{95}$ |
|---|---|---|---|---|---|---|---|---|---|
| MIQ-RS | Visual Subscale | 39.65 | 5.35 | 39.83 | 5.89 | 0.62 | [0.48, 0.72] | 3.48 | 8.10 |
| | Kinesthetic Subscale | 35.17 | 8.05 | 35.25 | 8.41 | 0.82 | [0.75, 0.88] | 3.48 | 8.10 |
| iFST | Raw difference (seconds) | 1.01 | 1.14 | 0.86 | 0.91 | 0.85 | [0.78, 0.9] | 0.39 | 0.91 |
| | Relative difference (% of error) | 25.03 | 21.73 | 25.00 | 23.45 | 0.82 | [0.74, 0.87] | 9.64 | 22.43 |
| | Constraint score (ratio difference) | 0.30 | 0.30 | 0.22 | 0.24 | 0.39 | [0.21, 0.54] | 0.21 | 0.50 |
| HLJT | Accuracy (%) | 0.90 | 0.07 | 0.92 | 0.07 | 0.76 | [0.59, 0.85] | 0.04 | 0.08 |
| | Reaction time (ms) | 1213.26 | 318.62 | 1086.12 | 302.02 | 0.80 | [0.44, 0.91] | 137.81 | 320.56 |
| | Biomechanical constraints (ms) | -233.96 | 186.55 | -212.06 | 194.64 | 0.71 | [0.6, 0.8] | 102.33 | 238.04 |

Abbreviations: HLJT: Hand Laterality Judgement Task; iFST: Imagined Finger Sequence Task; ICC: Intraclass Correlation Coefficient; $MDC_{95}$: Minimal Detectable Change (95% level); MIQ-RS: Movement Imagery Questionnaire-Revised Second edition; SD: Standard Deviation; SEM: Standard Error of Measurement.

### Results of the self-assessment questions

#### MIQ-RS

Regarding the sensory modality preference (visual or kinesthetic), the distribution of responses showed a majority of participants preferring a visual modality or having no preference compared to preferring a kinesthetic modality (Fig. S8a). Regarding the visual perspective preference (first-person or third-person), the distribution of responses showed a majority of participants preferring a first-person perspective or having no preference, compared to preferring a third-person perspective (Fig. S8b).

Regarding the use of kinesthetic imagery while performing visual items, the distribution of responses showed a majority of participants showing no use or moderate use (Fig. S8c). Regarding the use of visual imagery while performing kinesthetic items, the distribution of response showing a majority of participants showing moderate or high use (Fig. S8d).

**Supplementary Figure 8.** Boxplots and densities for the responses on self-assessment questions of the Movement Imagery Questionnaire-Revised Second edition (MIQ-RS). All questions were assessed on 11-point scales (0-10 points).

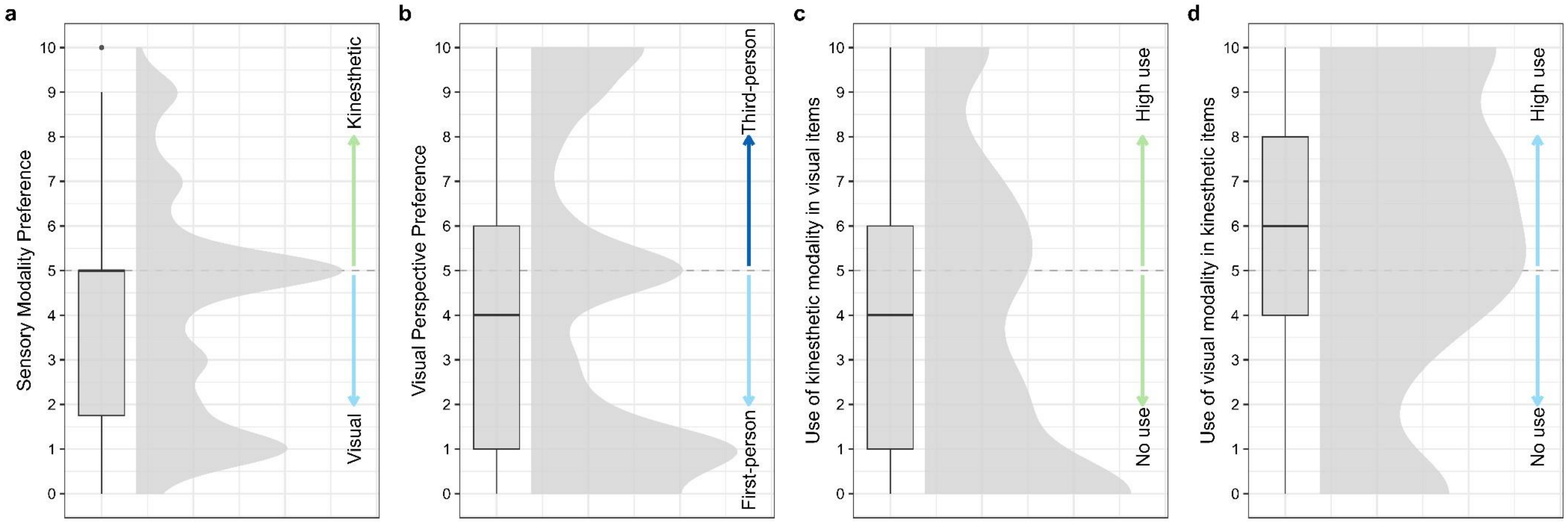

### iFST

Regarding the overall vividness, no differences between sequences could be observed based on the distribution of responses, with the majority of participants reporting moderate to high vividness (Fig. S9a). Regarding overall accuracy, a slight lower self-reported accuracy could be observed for the complex sequence, but both distributions show very limited self-perceived errors (Fig. S9b).

Regarding the use of the visual and kinesthetic modalities, no difference between sequences could be observed and the distributions show moderate to high use (Fig. S9c and Fig. S9d, respectively). Regarding the use of the first-person visual perspective, no difference between sequences could be observed and the distributions show high to very high use (Fig. S10e). Regarding the use of the third-person visual perspective, no difference between sequences could be observed and the distributions show low or very low use (Fig. S10f).

**Supplementary Figure 9.** Boxplots and densities for the responses on self-assessment questions of the Imagined Finger Sequence Task (iFST). All questions were assessed on 11-point scales (0-10 points).

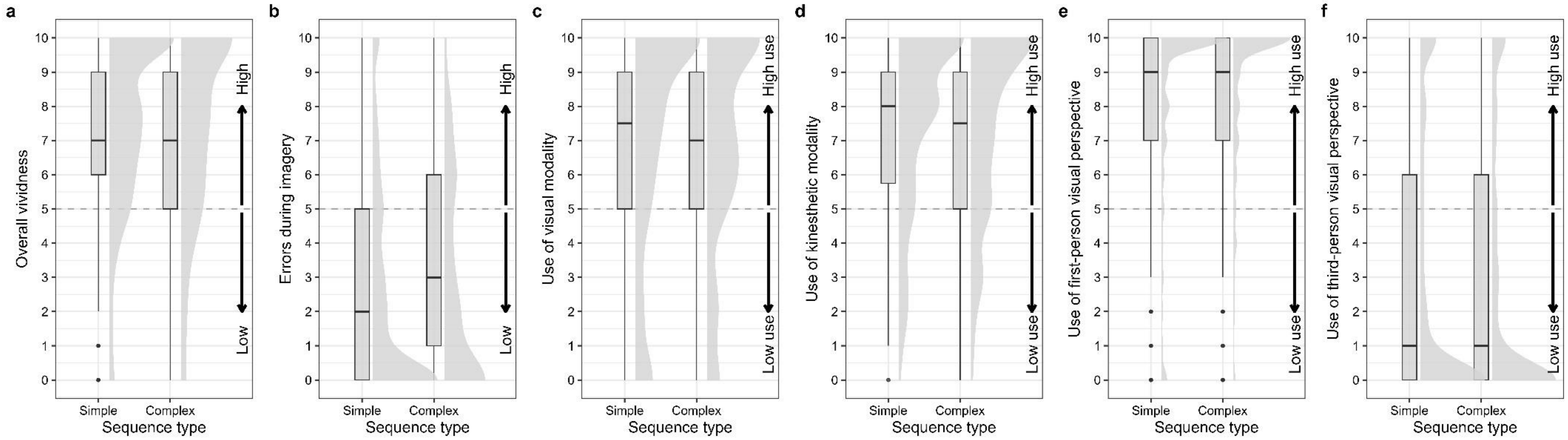

HLJT

Regarding the proportion of participants reporting having thought about their own hands, the majority of participants reported they did think about their hands, regardless of having being told about the concept of movement imagery or not (Fig. S10a). The proportion of participants reporting using imagery was slightly lower, but again the majority of participants reported they did use imagery, regardless of being aware or not about it (Fig. S11b). Those who used imagery primarily imagined both hands, as opposed to imagining only the left or right hand exclusively (Fig. S10c).

Regarding the use of the visual modality, the distribution showed participants reported high to very high use (Fig. S11a). Regarding the use of the the kinesthetic modality, the distribution reported low or no use, moderate use or high use (Fig. S11b). Regarding the use of the first-person visual perspective, the distribution showed participants reported high to very high use (Fig. S11c). Regarding the use of the third-person visual perspective, the distribution showed participants reported no or very low use (Fig. S11d).

**Supplementary Figure 10.** Proportion of participants according to their self-reported assessments in the Hand Laterality Judgement Task (HLJT).

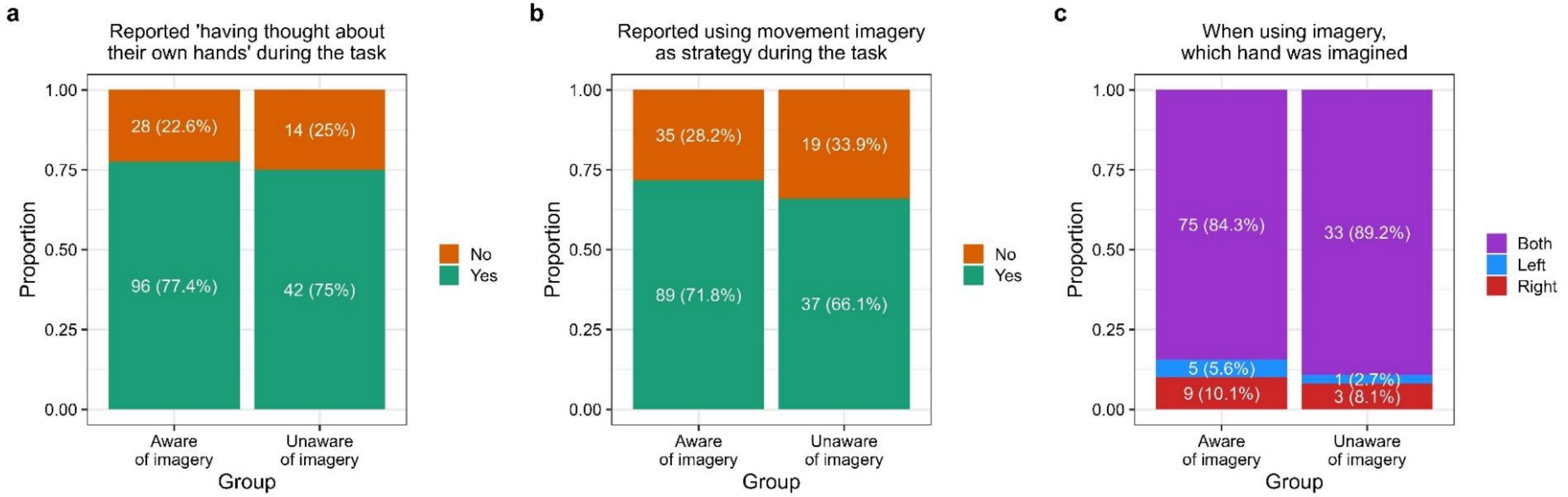


**Supplementary Figure 11.** Boxplots and densities for the responses on self-assessment questions of the Hand Laterality Judgement Task (HLJT). All questions were assessed on 11-point scales (0-10 points).

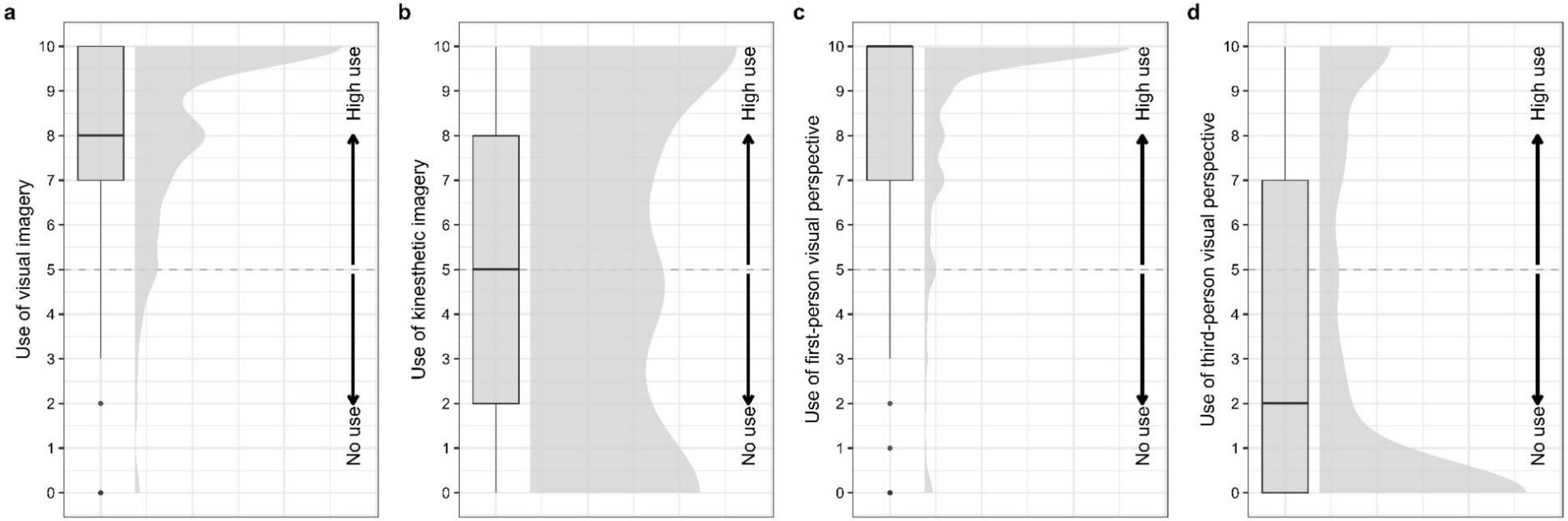
